# Deep homology and design of proteasome chaperone proteins in *Candida auris*

**DOI:** 10.1101/2025.05.14.654010

**Authors:** Jackson R. Rapala, Mohammad Siddiq, Patricia J. Wittkopp, Matthew J. O’Meara, Teresa R. O’Meara

## Abstract

A central tenet of biology is that protein structure mediates the sequence-function relationship. Recently, there has been excitement about the promise of advances in protein structure modeling to generate hypotheses about sequence-structure-function relationships based on successes with controlled benchmarks. Here, we leverage structural similarity to identify rapidly evolving proteasome assembly chaperones and characterize their function in the emerging fungal pathogen *Candida auris*. Despite the large sequence divergence, we demonstrate conservation of structure and function across hundreds of millions of years of evolution, representing a case of rapid neutral evolution. Using the functional constraints on structure from these naturally evolved sequences, we prospectively designed *de novo* chaperones and demonstrate that these artificial proteins can rescue complex biological processes in the context of the whole cell.

## Main Text

The causal relationship between sequence, structure, and function is central to our ability to assign function for newly discovered proteins and explore evolutionary relatedness based on inferred homology. While the importance of BLAST and other sequence similarity searches cannot be overstated, these approaches can fail to detect evolutionarily related proteins with high levels of sequence divergence (*1*). However, since structure mediates the sequence-function relationship, signatures of shared evolutionary history are often better preserved at the level of structure than sequence (*2*, *3*). The recent advances in computational protein structure prediction have enabled large-scale comparative structural analyses of proteins across broad swathes of evolution (*3–7*). In turn, it has become possible to experimentally probe the constraints and flexibilities within which sequence-structure relationships evolve (*8–10*). Further, multiple protein sequences—homologous or convergently evolved—often result in the same or similar structure and thus similar function (Figure 1A). Through this understanding of the structural requirements for protein function, it should then be possible to generate *de novo* designed proteins that are disconnected from evolutionary history and are yet still able to fulfill the biological requirements of the cell (Figure 1A) (*11*).

**Fig. 1.**
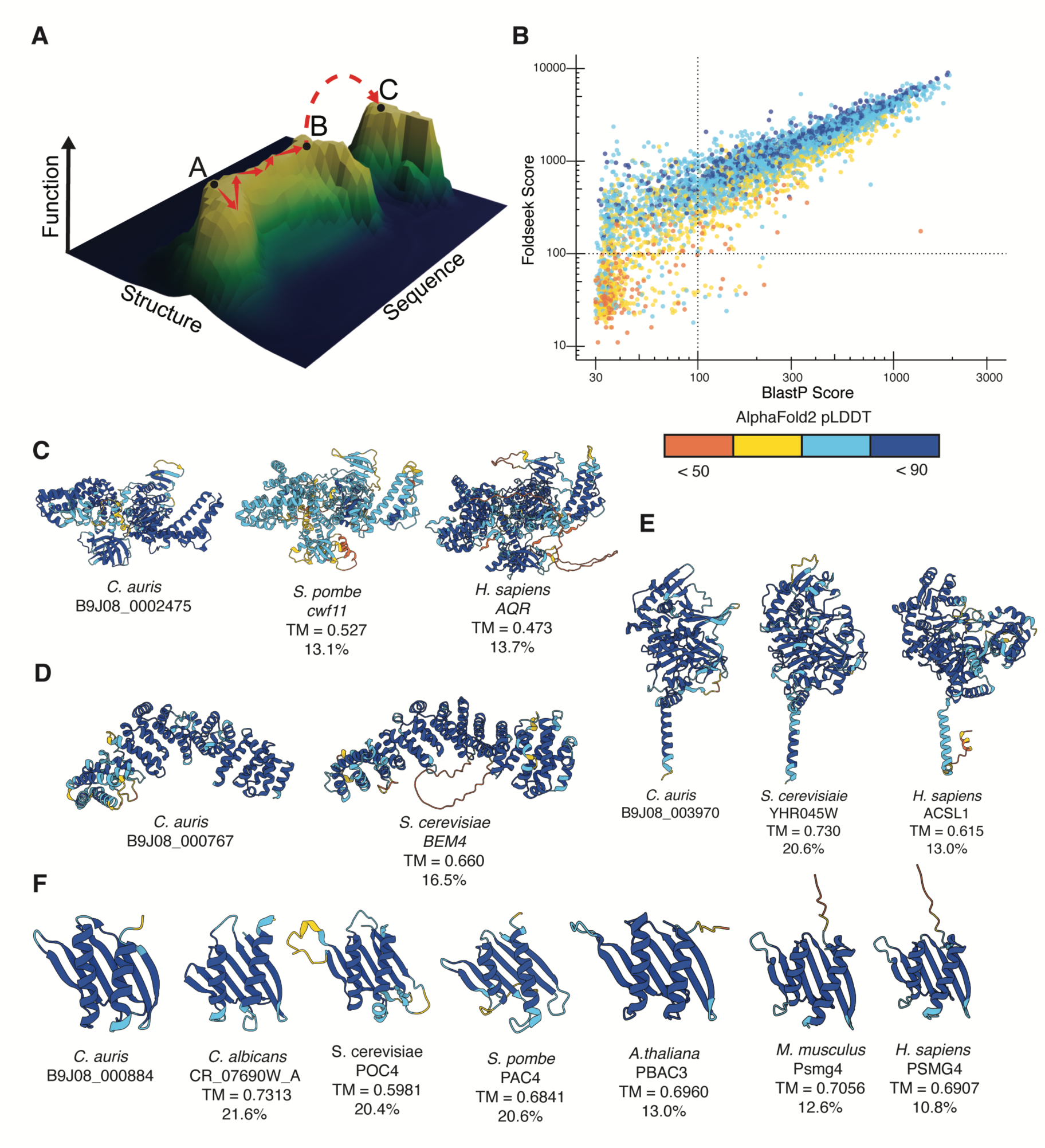
Structural homology can be used to predict protein function in the absence of sequence homology. **(A)** Diagram of a hypothetical fitness landscape, with the X axis as sequence, Z axis as structure, and Y axis as fitness. **(B)** Detection of remote homology of *C. auris* proteins by structure (Foldseek with AlphaFold Database predicted structures) vs. sequence (BlastP). For each of the 5,008 proteins for which there is an AFDB structure, a plot of the best bit-scores to a protein in a Swiss-Prot proteome colored by the pLDDT, where the 100-bit threshold is indicated. AlphaFold 2 models of *C. auris* proteins and structure matches identified by Foldseek for B9J08_002475 **(C)**, B9J08_000767 **(D)**, B9J08_0003970 **(E)**, and B9J08_000884 (Poc4) **(F)**. Models are colored by pLDDT by residue. Percentage indicates amino acid identity to the *C. auris* protein.

Here, we explicitly tested the model that vastly divergent protein-coding sequences predicted to encode similar structures result in similar functional phenotypes across evolutionary time. These represent cases of neutral evolution, where sequence changes do not result in the structural changes that can then be selected for or against. In the non-model human fungal pathogen *Candida auris,* we identified rapidly evolving but putatively structurally conserved proteasome assembly chaperones and determined that they have conserved roles in proteasome function and thermotolerance. The rate of evolution of these assembly chaperone proteins differed even amongst interacting partners, suggesting our understanding of co-evolutionary rates based on sequence divergence might be biased by those with clear membership in defined orthogroups. We used deep learning to explore protein sequence space and designed artificial proteasome chaperones to maintain a conserved structure and rescue mutant phenotypes. Our work demonstrates how modern computational methods, in combination with experimental characterization, can be used to explore the evolution of sequence-structure-function relationships and, from the insights gained, enable us to create functional molecules released from the constraints of evolutionary history.

### Structural homology in the absence of sequence homology predicts protein function

While structure-based similarity search outperforms sequence-based homology in retrospective benchmarks, it remains challenging to assess the functional role of these predictions (*12*, *13*). Signatures of homology are better preserved at the level of structure, therefore structure-based homology search should enable better inferences of gene functions across large evolutionary distances where sequence-based methods fail. To test this prediction, we compared the predictive power of Foldseek, a structure-based homology search algorithm, to BlastP, a sequence-based homology search algorithm, to detect orthologs for *C. auris* proteins amongst the proteins in the Swiss-Prot database (Figure 1B, Supplemental Figure 1) and observed a high correlation between their respective bit-scores over the 5,008 genes that had predicted AlphaFold2 structures (Spearman Rank Correlation: 0.932, p-value < 0.0001). From this analysis, we identified 204 genes with significant Foldseek E-values, defined as less than 1e-5, and insignificant BlastP E-values representing proteins where structural homology may be able to predict function in the absence of sequence homology.

For example, the protein B9J08_002475 and its sequence orthologs are *Metschnikowiaceae-*lineage restricted (Figure 1C) but have significant structural homology with the Aquarius family of RNA helicases that function in the U2 snRNP spliceosome (Supplemental Table 5) despite sharing less than 15% sequence identity. Similarly, B9J08_000767, which is a *Candida*-lineage restricted protein, has significant structural similarity to the Bem4 guanidine exchange factor (GEF) in *S. cerevisiae* but only 16.5% sequence identity (Figure 1D, Supplemental Table 6). We are also able to leverage Foldseek against uncharacterized proteins present broadly throughout fungi, including B9J08_003970, which has structural similarity with mammalian long-chain fatty acid acyl-CoA synthetases 1, 5 and 6 (Figure 1E, Supplemental Table 7).

Selection acts not just on single proteins but the entire functional unit of a protein complex (*14*, *15*); therefore, we were particularly interested in the case of two proteins putatively involved in the highly coordinated process of proteasome assembly. We identified the uncharacterized protein B9J08_000884, with a similar structure to the Poc4 assembly chaperone protein, and B9J08_003917, which is annotated as a homolog of the Irc25 proteasome assembly chaperone (Supplemental Table 8) (*16–18*). We will refer to these two proteins as Poc4 and Irc25, following nomenclature from *S. cerevisiae*. Proteins with structural similarity to Poc4 were widely present throughout Eukarya, and despite sequence identities below 20%, the overall topology of the protein domains is predicted to be conserved (Figure 1F, Supplemental Figure 2A). Given the overall high sequence and structure conservation of the eukaryotic proteasome, the sequence divergence in the assembly chaperones was not anticipated, as proteins involved in related processes are expected to co-evolve at similar rates (*14*, *15*).

### Sequence divergence in proteasome assembly chaperones

To better understand how distinct components of the proteosome might have coevolved, we used amino acid and syntenic analysis to search for homologs of proteosome assembly components across fungi. The putative orthologs of Poc4 and Irc25 function as a heterodimer and mediate assembly of α-subunits into the heptameric ring structure of the proteosome, interacting especially with the α5-subunit (Pup2), before dissociating during β-subunit assembly (Figure 2A) (*19*). The Pba1-Add66 complex also contributes to α-ring assembly around the Ump1 chaperone. We found that while Pup2 proteasome subunit and the Ump1 assembly chaperone had detectable homologs throughout *Saccharomycotina*, the CTG clade lacked a clearly identifiable sequence that resembled the Poc4 proteins found in other yeast, and putative homologs of the other proteasome chaperones were at times only identifiable because of synteny (Figure 2B).

**Fig. 2.**
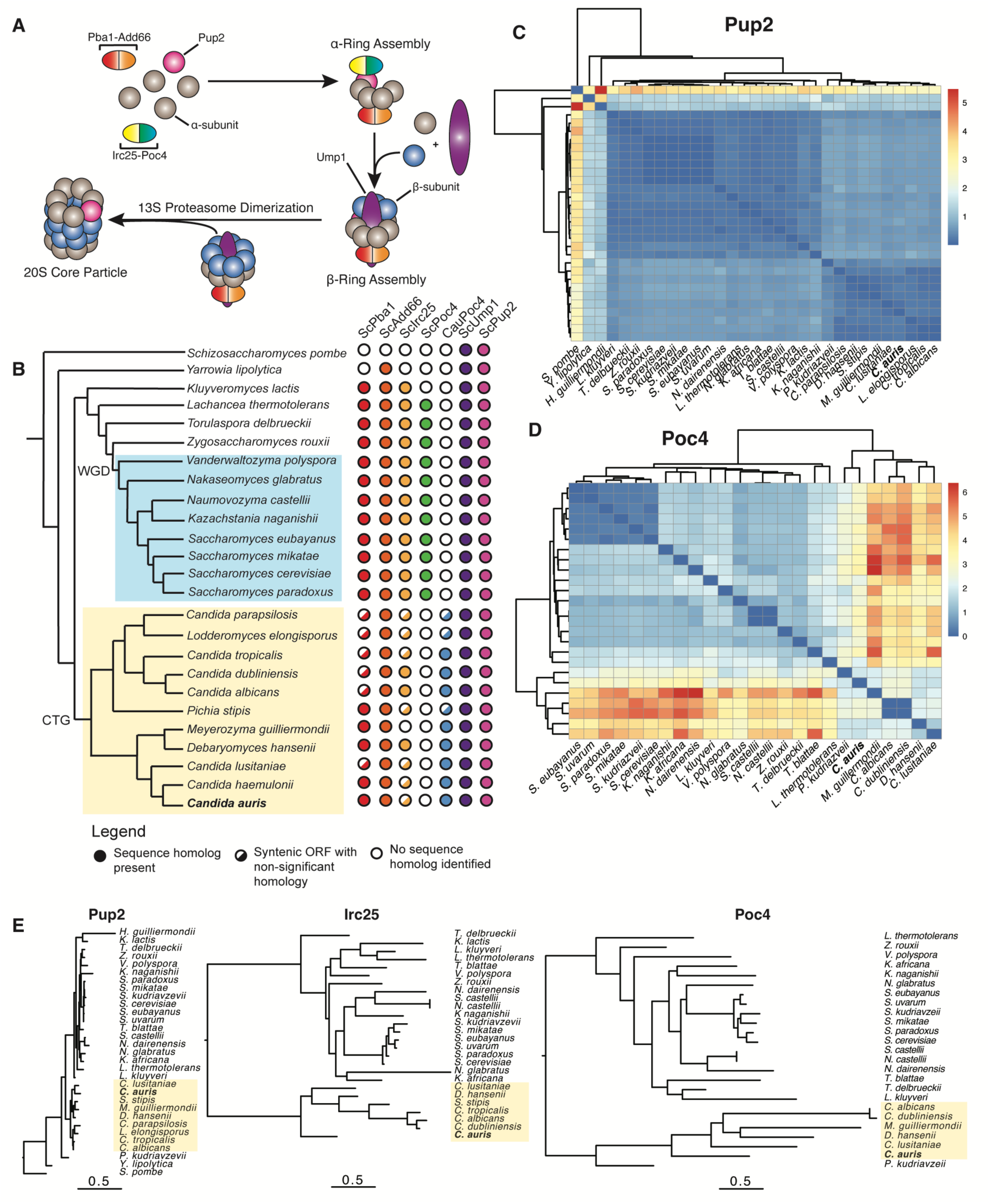
*C. auris* Poc4 is sequence divergent, but not structurally divergent, from known Poc4 assembly chaperones. **(A)** Schematic of 20S core particle assembly. The Pba1-Add66 and Irc25-Poc4 complexes initiate assembly of α-subunits into a heptameric ring structure, where Irc25-Poc4 directly interact with the Pup2 subunit. Before β-subunit incorporation, Irc25-Poc4 dissociates and Ump1 enters the central cavity to create a half proteasome. Two half proteasomes dimerize to create a 20S core particle. **(B)** Evolutionary relationship of analyzed yeast species shows rapid loss of sequence identity among proteasome assembly chaperones. Sequences with similarity to the indicated *S. cerevisiae* and *C. auris* proteins were identified using BlastP from NCBI using an E-value cutoff of 10^-5^ and requiring hits to reciprocal best matches. CTG indicates the clade where the CUG codon is translated as serine instead of leucine. WGD denotes the whole genome duplication clade. Pairwise comparisons of maximum likelihood distances between Pup2 **(C)** and Poc4 **(D)** sequences. **(E)** Evolutionary rate of Pup2, Irc25, and Poc4 sequences among indicated yeast species identifies rapid change in chaperone proteins. Scale = 0.5 substitutions per site.

The variable presence of different proteosome components suggested that the folding chaperones and core components may have different evolutionary rates from each other. To compare the extent of divergence among core proteins and folding chaperones, we used the putatively orthologous sequences from BlastP and inferred the maximum likelihood phylogeny describing the evolutionary history of each gene using the best-fit model of sequence evolution, as determined by Akaike information criterion (Supplemental Table 9, Supplemental Figure 3A-C). The proteins investigated displayed vastly different rates of divergence. The evolutionary distance between Pup2 orthologs, for example, is much smaller than the difference between chaperone orthologs (Fig. 2C, E). This difference in rate of evolution among the chaperone proteins is not unique to any species lineage but is true across all taxa (Fig. 2D, E). Interestingly, Poc4 appears to be evolving faster than its co-chaperone Irc25 (Figure 2E, Supplemental Figure 3D), which would not be expected for proteins in complex without other known functions. One possible reason for this discrepancy in rate is expression levels, as higher expression is associated with lower evolutionary rate (*20*). Previously published RNAseq data show that Poc4 expression is much lower than other core particle assembly chaperones (Supplemental Figure 3E) (*21*).

### *C. auris POC4* encodes a *bona fide* proteasome assembly chaperone

While we observed structural similarity to known Poc4 proteasome assembly chaperones, the extreme sequence divergence suggested that its role in proteasome assembly might not be preserved. To test whether *C. auris* Poc4 retained canonical interactions with the proteasome, we performed co-immunoprecipitation assays with an ALFA-tagged variant of Poc4 and the α-subunits of the 20S core particle. We found this physical interaction was preserved (Figure 3A and Supplemental Figure 4A-B), suggesting that the capacity to act as an assembly chaperone had persisted in Poc4 even as its amino acid sequence changed beyond recognition.

**Fig. 3.**
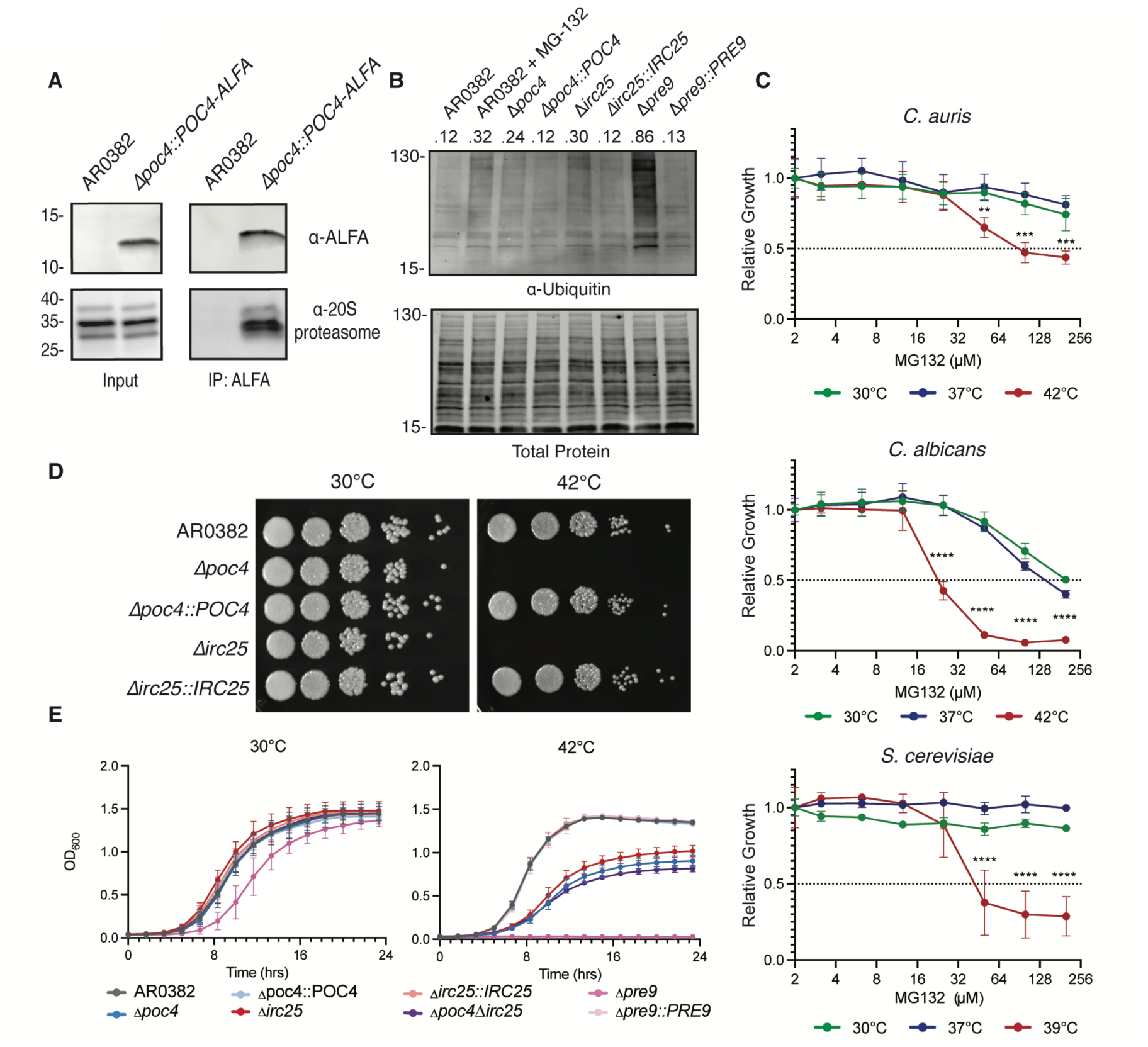
*C. auris POC4* encodes a *bona fide* proteasome assembly chaperone. **(A)** *C. auris* Poc4 physically interacts with the 20S proteasome. Immunoprecipitation of Poc4-ALFA with anti-ALFA co-purifies proteasome α-subunits, which do not co-purify with untagged controls. Numbers on left of blots indicate approximate size in kDa. **(B)** Total protein ubiquitination increases in *C. auris* upon loss of *POC4* or *IRC25*. The indicated strains were subcultured into fresh media for 6 hours at 30 °C before protein extraction and western blotting. Numbers above lanes indicate relative Ubiquitin intensity normalized to total protein intensity. **(C)** Proteasome inhibition reduces fungal growth at high temperatures. Dose-response curves of *C. auris*, *C. albicans,* and *S. cerevisiae* to MG-132 at the indicated temperatures. OD_600_ at each drug concentration is normalized to the growth of each strain in the absence of drug to determine relative growth. Data show mean ± standard deviation across 3 biological replicates. **(D)** Poc4 and Irc25 are both required for growth at 42°C. Representative spot plates of *C. auris* grown at 30 °C and 42 °C. Indicated strains were diluted 10-fold and plated onto YPD agar and incubated for 48 hours before imaging. **(E)** Growth curves of indicated *C. auris* strains at 30 °C and 42 °C. Growth was measured by OD_600_ every 10 minutes for 24 hours. Data shown are the mean ± standard deviation across 4 biological replicates every 80 minutes for clarity. Statistical significance among different groups was calculated using one-way ANOVA, with Dunnett’s post hoc tests for multiple comparisons. *p ≤ 0.05; **p ≤ 0.01; ***p ≤ 0.001; n.s., p > 0.05.

Next, we directly characterized whether this physical interaction was involved in the same cellular functions. Disruption of proteasome function through genetic perturbations (e.g., deletion of Pre9 subunit) or pharmacological inhibition (e.g., MG-132 treatment) increased the level of ubiquitinated proteins within a cell (Figure 3B). We reasoned that disruption of Poc4 should also produce these phenotypes. Consistent with our model, we observed increased total ubiquitinated proteins in Poc4 and Irc25 deletion strains relative to the wild-type AR0382 strain (Figure 3B).

MG-132 had limited effects on fungal growth at permissive temperatures but significantly reduced growth at high temperatures across three fungal species (Figure 3C), demonstrating a conserved connection between proteasome inhibition and thermotolerance across organisms. Similar results were observed with a 2^nd^ proteasome inhibitor, Bortezomib, but not general antifungal drugs (Supplemental Figure 5A-B). Furthermore, *C. auris* cells lacking Poc4 or Irc25 are hypersensitive to MG-132 even at permissive temperatures (Supplemental Figure 5C). Consistent with the pharmacological data, loss of either *IRC25* or *POC4* resulted in a temperature sensitive phenotype, where both mutants are unable to grow at 42°C, but growth is restored upon complementation (Figure 3D). Furthermore, the *Δirc25* and *Δpoc4* individual deletion strains phenocopy each other, but deletion of both genes does not alter the phenotype beyond a single deletion, suggesting non-redundant functions for these proteins (Figure 3E). By contrast, deletion of the gene encoding Pre9, a proteasome subunit, results in temperature sensitive phenotype even at 37°C (Supplemental Figure 6A-B). Taken together, these experimental data demonstrate that, despite losing detectability as a sequence homolog *C. auris* Poc4 has retained a conserved evolutionary role in proteasome function.

### *S. cerevisiae* Poc4 can function in *C. auris,* despite considerable sequence divergence

The *C. auris* Poc4 role in proper proteasome function, despite its highly divergent sequence, suggests considerable flexibility in sequence-structure relationships within which evolution can operate while preserving essential functions. Recent advances in prediction of multi-subunit protein complexes have made it possible to assess if the physical interactions are preserved. We used AlphaFold 3 to fold the proteasome core particle during assembly with both the Pba1-Add66 and Irc25-Poc4 chaperone complexes, starting with the *S. cerevisiae* complex as a control. We observed high predicted structural quality and similarity with the experimentally characterized complex obtained from cryo-electron microscopy (*22–24*) (Supplemental Figure 7A-B, ipTM = 0.78). Using the same parameters, we folded the *C. auris* proteasome complex and observed structural similarity with the *S. cerevisiae* proteasome complex (Figure 4A, Supplemental Figure 7C, ipTM = 0.81). Comparing the *S. cerevisiae* and *C. auris* complexes, we observe conserved position of Poc4 above the pore of the proteasome. Notably, in addition to the expected predicted hydrogen bonds between the Irc25 and Poc4 protein along the dimerization interface, we observed 5 predicted hydrogen bonds between Poc4 and Pup2, suggesting that Poc4-proteasome interactions are crucial for chaperone activity (Figure 4B, Supplemental Figure 7B,C). Of the three residues in *S. cerevisiae* Poc4 that are required for Poc4-Pup2 interactions (*25*) none are conserved in *C. auris* Poc4, and two of these differing residues have major changes in amino acid chemistry (Supplemental Figure 7D). Given these differences, we next sought to characterize whether the Poc4 homologs were functionally interchangeable.

**Fig. 4.**
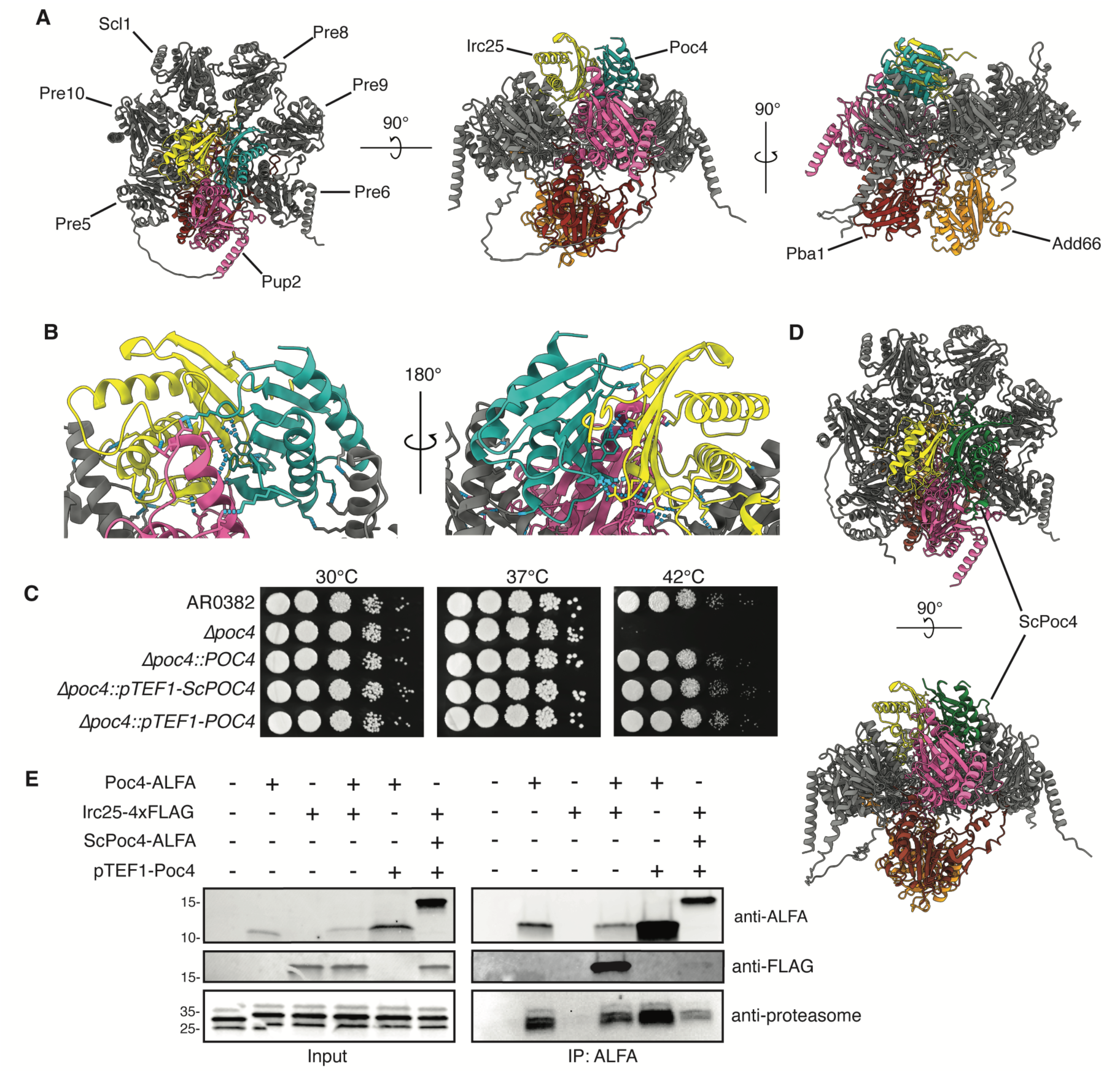
*S. cerevisiae* Poc4 can function in *C. auris,* despite sequence divergence. **(A**) AlphaFold 3 model of *C. auris* proteasome subunits during core particle assembly. **(B)** Close up view of Poc4-Irc25-Pup2 interface from structure in (A). Predicted hydrogen bonds indicated by dashed blue lines. **(C)** *S. cerevisiae* Poc4 is sufficient for growth at 42 °C in *C. auris*. Representative spot plates of indicated *C. auris* strains grown at 30 °C, 37 °C, and 42 °C. Indicated strains were diluted 10-fold, plated onto YPD agar, and incubated for 48 hours before imaging. **(D)** AlphaFold 3 model of *C. auris* proteasome, with *S. cerevisiae* Poc4 replacing *C. auris* Poc4 **(E)** *S. cerevisiae* Poc4 physically interacts with the 20S proteasome and Irc25. Co-immunoprecipitation of Irc25 and proteasome α-subunits with *C. auris* and *S. cerevisiae* Poc4 constructs. Numbers on left indicate approximate size in kDa. Images and blots are representative of at least three biological replicates.

The general conservation of the modeled protein complexes suggests that, despite sharing only 20% sequence identity, the Poc4 proteins may be functionally interchangeable. However, the observed differences at key residues may create structural incompatibilities if a Poc4 variant is placed in context of heterologous proteasome components. To evaluate the extent of functional conservation on Poc4 chaperones, we tested if a CUG-optimized Poc4 allele from *S. cerevisiae* would complement the temperature-sensitive growth defects of the *C. auris* deletion strain (Figure 4C). We expressed *ScPoc4* under a constitutive promoter in the *C. auris Δpoc4* background and observed full restoration of thermotolerance, suggesting that proteasome assembly activity is being rescued. Complementation of *Δpoc4* with a promoter-matched *C. auris* Poc4 also restores growth at 42 °C, but growth was not enhanced over that of the respective wildtype. Of note, we were unable to complement with the *S. cerevisiae* Poc4 allele under control of the endogenous *C. auris* Poc4 promoter because this strain failed to express at 42 °C (Supplemental Figure 8A-C).

To understand the mechanism of rescue, we predicted the fold of the *C. auris* proteasome complex as above but replaced the endogenous Poc4 with *S. cerevisiae* Poc4 using AlphaFold 3 (Figure 4D, Supplemental Figure 7E). Despite considerable sequence divergence, *S. cerevisiae* Poc4 is still predicted to interact with *C. auris* Irc25 and the α5 subunit with DockQ2 scores of 0.42 and 0.25, respectively. This suggested that although specific interaction residues were disrupted, the overall interaction interface is maintained. To test this experimentally, we immunoprecipitated the Poc4 and found that in *C. auris, S. cerevisiae* Poc4 physically interacts with the proteasome α-subunits and Irc25 (Figure 4E, Supplemental Figure 4C-D), albeit to a lesser degree. Thus, we show a deeply conserved role of Poc4 across fungi that has persisted despite erosion of sequence signatures of homology.

### Prospective testing of designed chaperone proteins

Our results suggest that certain structural features are sufficient for Poc4 proteasome assembly functions, but the generality of this inference is limited by the fact that Poc4 variants from the two species have a shared evolutionary history. Therefore, we wanted to test whether an artificially designed protein would be sufficient to rescue the mutant phenotype. We used Frame2Seq (*26*), a state-of-the-art method for conditional sequence generation, to design novel sequences for the predicted native *C. auris* Poc4 backbone geometry, conditional on the structural fold of the native Poc4 and predicted interactions with Irc25. To maintain the interaction with Irc25, we fixed the 29 residues at the predicted interface and excluded cysteine residues. We wanted to achieve a level of sequence diversity comparable to natural variation, which we were able to attain using the default temperature parameters (Supplemental Figure 9).

We then assessed how these generated sequences related to native sequences using the ESM2 model to embed the native Poc4 and Irc25 sequences, the orthologous sequences identified through Foldseek, the POC3_POC4 pfam family (PF10448), and the re-designed Poc4 chaperone sequences into a continuous high-dimensional latent sequence space and used UMAP to project the embedded representations to 2-dimensions. As expected, we observed moderate clustering among the Poc4 homologs, Irc25 homologs, and designed sequences, but some mixing between the clusters (Figure 5A). This gave us confidence that our designs were, at least at the sequence level, distinct from natural Poc4 chaperones. Since our aim was to test the structural basis for function, we selected 5,000 Frame2Seq designs, used AlphaFold2 to re-fold each sequence, and compared the pLDDT with the Frame2Seq score and sequence conservation (Supplemental Figure 9). For the 82 variable residues, recovery of the native Poc4 sequence ranged between 50-70 percent identity and 98.9% had an average pLDDT score > 90, suggesting the scaffold is designable by Frame2Seq. From these, we selected 5 designs for experimental testing, with 61 to 64% sequence identity to *C. auris* Poc4 but without homology by BlastP to other proteins. We found that all designed proteins were expressed in the *Δpoc4* deletion background, though only Seq2523 showed significant rescue of thermotolerance at 42 °C (Figure 5B-C, Supplemental Figure 10A-B).

**Fig. 5.**
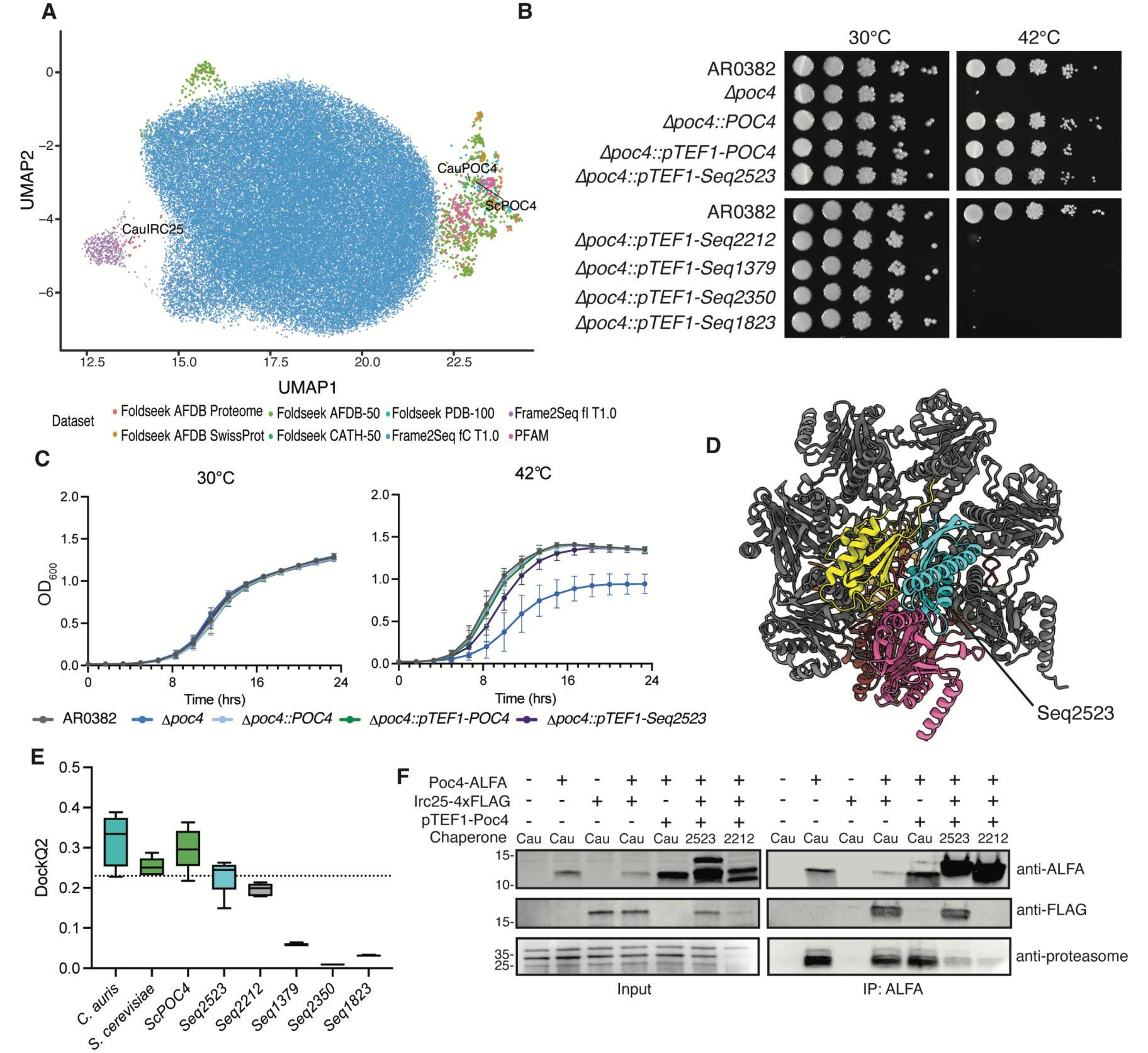
*de novo* Poc4 chaperone designs are functional *in vivo.* **(A**) Embedding of Poc4 and Irc25 sequence space, with both natural and designed protein sequences. **(B)** Seq2523 rescues *C. auris* growth at 42 °C. Representative spot plates of the indicated *C. auris* strains grown at 30 °C and 42 °C. Indicated strains were diluted 10-fold, plated onto YPD agar, and incubated for 48 hours before imaging. **(C)** Growth curves of indicated *C. auris* strains at 30 °C and 42 °C. Growth was measured by OD_600_ every 10 minutes for 24 hours. Data shown is the mean ± standard deviation across 4 biological replicates every 80 minutes for clarity. **(D)** AlphaFold 3 model of *C. auris* proteasome subunits during core particle assembly, with designed chaperone Seq2523 instead of the native Poc4. **(E)** Seq2523 is predicted to dock better with the proteasome than other tested designs. DockQ2 scores between Poc4 and Pup2 in the indicated structures. Data shown is the mean ± standard deviation across 5 models. Structures with activity are colored in green *(S. cerevisiae*) and blue (*C. auris*), while non-active structures are in gray. Dashed line indicates significance cutoff. **(F)** Co-immunoprecipitation of Irc25 and proteasome α-subunits with designed chaperone proteins. Numbers on left indicate approximate size in kDa. Images and blots are representative of three biological replicates.

That any, but not all, designs were able to rescue *Δpoc4* suggested that the structural features substantially contribute to but do not fully explain the functional constraints on Poc4. During the design, we selected for interaction with Irc25 but not the rest of the proteasome. To retrospectively test whether this difference in interaction would explain the variable ability to rescue, we used AlphaFold3 to predict the proteasome assembly complex, substituting each design for the wildtype Poc4 (Figure 5D, ipTM = 0.81). We observed similar predicted structures as the wildtype interactions with the α-subunits (Supplemental Figure 10C). We then calculated DockQ2 scores between all interacting subunits of the complex for all structures (ipTM range 0.77-0.81). Strikingly, only the functional design (Seq2523) had DockQ2 scores with both Irc25 and Pup2 over the significance threshold (Figure 5E) (*27*, *28*). This data is further supported by immunoprecipitation, where we observed stronger associations with both Irc25 and the proteasome α-subunits with the functional Seq2523 design than the non-functional Seq2212 design (Figure 5F).

## Discussion

Over the past decade, advances in genome sequencing have drastically increased the number of available gene/protein sequences, and our ability to compare and define relationships between sequences is critical for our understanding of genome function, evolution, and biotechnology. Central to this process are sequence-based search algorithms, such as BLAST, which have revolutionized our understanding of comparative genomics and evolution. However, these methods often fail to identify true homologs once sequence identity falls below 30%, reaching the so-called twilight zone (*1*). This issue has widespread effects: recent estimates suggest that 40 to 45% of all lineage-restricted genes in fungi are due to homology detection failure and not *de novo* origination (*29*, *30*). Consequently, homologous proteins that evolve rapidly in sequence have been excluded from comparative studies on protein evolution. Here, we leveraged comparisons of protein structure to overcome the limitations of sequence analysis and identify cases of structural homology in the emergent fungal pathogen *C. auris*. When focusing on proteins within the BlastP twilight zone, we identified a hypothetical protein with structural similarity to the Poc4 assembly chaperone in *S. cerevisiae*. We found that yeast proteasome assembly chaperones rapidly evolve at rates that would obfuscate homology detection by sequenced-based approaches. Yet, despite the sequence divergence between *C. auris* Poc4 and *S. cerevisiae* Poc4, we found that the orthologs have evolutionarily conserved function in proteasome assembly and are functionally interchangeable in *C. auris*.

Our initial observation that proteasome assembly chaperones are rapidly evolving was surprising because the client proteins of these chaperones, the subunits of the proteasome, are highly conserved in sequence and structure throughout Eukarya (*31*, *32*). While the sequence divergence and functional redundancy between human and yeast proteasome assembly chaperones were observed during their initial discovery, the rapid divergence and evolutionary implications were not explored (*16*, *17*). We had expected that proteins that physically interact and codetermine function would also have similar coevolutionary rates (*14*). However, our observations suggest this does not have to be the case; rather, they support the idea that in any given system, a wide range of tolerable sequences may be sufficient for a given structure (*33*). In the absence of a change in structure, there is then nothing for selection to act upon, thus resulting in the accumulation of sequence changes over time. Furthermore, as the Poc4 sequence diverges in a lineage, epistatic interactions between and within chaperone proteins and proteasome subunits may further entrench sequence changes, preventing any sort of reversion back to an ancestral sequence (*34–37*).

Protein evolution is strongly shaped by historical contingency, meaning the paths available to and taken by evolution are strongly affected by prior evolutionary events (*36*, *37*). A consequence of this contingency is that many parts of protein sequence space likely exist that have not been sampled over evolution. To explore the unsampled or otherwise evolutionarily inaccessible protein space, we leverage recent advances in protein design and prospective testing to evaluate the sufficiency of a specific fold to fulfill a function required for cellular survival. This approach expands the sequence space of functional proteins beyond what can be achieved through deep mutational scanning or natural and lab evolution of new traits and functions, as these single-step approaches can be severely limited by constraints imposed by epistatic interactions (*38–40*). In our approach, we were able to simultaneously mutate 40 of the 111 residues and still maintain function. Through total protein re-design, it is possible to explore epistatic relationships between amino acids and potentially access high fitness variants that are unlikely to emerge through stepwise evolutionary approaches.

In recent years, the scale and feasibility of protein design has greatly increased with our ability to predict protein structures from a sequence, synthesize them, and validate their functions (*6*, *7*, *41*, *42*). We and others have previously published examples of redesigned enzymes with higher efficiency or stability than their natural counterparts, with a range of medical and technical applications (*43–46*). Nevertheless, most designed proteins are generated for *in vitro* use, and few examples exist of such proteins being functional *in vivo* (*47*, *48*). A limitation of this work is that our designed sequences maintained some identity with the natural variant, instead of a 0% identity replacement. A strength of the study is that we did not require an experimentally characterized *C. auris* protein structure to perform the design. However, we recognize that our ability to design and test these chaperones is greatly facilitated by the small, single domain nature of the target protein and a simple phenotypic test for function. As proteins can vary greatly in size, domain, architecture, and function, this approach will not be feasible for all proteins. Additionally, a more detailed biochemical characterization of the designed proteins may further explain differences in activity. Despite this, we find that computational measures of protein docking are sensitive enough to identify potential differences between functional and non-functional designs. This observation, along with other studies that directly integrate diffusion and docking models into protein design workflows, suggests that modeling endogenous protein-protein interactions of designed sequences is a critical step in identifying protein designs with desired activity (*44*). Overall, our work provides an example where we can modulate each aspect of the sequence-structure-function relationship to understand the functional constraints on protein evolution.

## Author contributions

Conceptualization: JRR, MS, MJO, TRO

Methodology: JRR, MS, MJO, TRO

Investigation: JRR, MS, MJO, TRO

Visualization: JRR, MS, MJO, TRO

Funding acquisition: JRR, MS, PJW, MJO, TRO

Project administration: TRO, MJO

Supervision: PJW, TRO, MJO

Writing – original draft: JRR, MS, MJO, TRO

Writing – review & editing: JRR, MS, PJW, MJO, TRO

## Data and materials availability

All data, code, and materials used in the analysis is available in the text or on Github (https://github.com/maomlab/poc4). Strain used in this will be sent following standard materials transfer agreements (MTAs).

## Supplementary Materials

Materials and Methods

Supplementary Text

Figs. S1 to S10

Tables S1 to S10

## Materials and Methods

### Strains and Culture Conditions

A list of all strains used in this study is included in Table S1. Clinical *C. auris* isolates were obtained through the CDC/FDA Antibiotic Resistance Isolate Bank. Unless specified, cells were cultured at 30°C in YPD liquid media (1% yeast extract, 2% peptone, 2% dextrose), with constant shaking. For solid medium, 2% agar was used. All strains were maintained as frozen stocks in 25% glycerol at -80°C.

### Genomic DNA Extraction

Genomic DNA was isolated using a PCA extraction method. Yeast cells were cultured overnight in YPD liquid at 30°C and harvested by centrifugation. Cells were resuspended in lysis buffer (2% (v/v) Triton X-100, 1% (w/v) SDS, 100mM NaCl, 10mM Tris-Cl, 1mM EDTA) and disrupted by bead beating. Released DNA was extracted into PCA and then into chloroform. The resulting DNA was then purified by ethanol precipitation and resuspended in water before treatment with RNase A. RNase was heat-inactivated and extracted DNA was further purified by ethanol precipitation and resuspended in water.

### Primers and Plasmids

A list of all plasmids used in this study is included in Table S2. A list of all primers used in this study is included in Table S3. Cassettes for transformation of *C. auris* were maintained in the multiple cloning site of the pUC19 cloning vector and assembled from fragments using the NeBuilder HIFI DNA Assembly Master Mix (NEB #E2621) or the Codex DNA Gibson Assembly Ultra Master Mix (Codex DNA #GA1200) according to the manufacturers’ instructions. All plasmid assemblies were confirmed by Nanopore sequencing.

### Plasmid and Strain Construction

**pTO301/CauTO467:** The plasmid backbone was amplified from pTO139 using oTO590 and oTO591. The NAT cassette was amplified from pTO137 using oTO847 and oTO668. The NAT cassette was flanked by 500bp of regions homologous to the regions immediately up and downstream of *POC4* (B9J08_000884) amplified from *C. auris* AR0382 genomic DNA using oTO1734-oTO1735 and oTO1736-oTO1737, respectively. The repair cassette was amplified using oTO18 and oTO19 and transformed into AR0382 to generate CauTO467 and CauTO468 from independent transformations.

**pTO306/CauTO497:** The plasmid backbone was amplified from pTO139 using oTO590 and oTO591. The ADH1 terminator and NEO cassette was amplified from pTO169 using oTO1066 and oTO668. The 500bp upstream and ORF of *POC4* (B9J08_000884) was amplified from *C. auris* genomic DNA using oTO1734 and oTO1842, while the downstream 500bp of homologous sequence was amplified from oTO1736 and oTO1737. The repair cassette was amplified using oTO18 and oTO19 and transformed into CauTO467 to generate CauTO497.

**pTO315/CauTO496:** The plasmid backbone was amplified from pTO139 using oTO590 and oTO591. The NAT cassette was amplified from pTO137 using oTO847 and oTO668. The NAT cassette was flanked by 500bp of regions homologous to the regions immediately up and downstream of *IRC25* (B9J08_003917) amplified from *C. auris* genomic DNA using oTO1928-oTO1929 and oTO1930-oTO1931, respectively. The repair cassette was amplified using oTO18 and oTO19 and transformed into AR0382 to generate CauTO496.

**pTO322/CauTO550:** The plasmid backbone was amplified from pTO139 using oTO590 and oTO591. The ADH1 terminator and NEO cassette was amplified from pTO169 using oTO1066 and oTO668. The NEO cassette was flanked upstream by the ORF of *IRC25* (B9J08_003917) and 500 bp of upstream sequence, which was amplified from *C. auris* genomic DNA using oTO1928 and oTO1980. A downstream flanking sequence was amplified from *C. auris* genomic DNA using oTO1930-oTO1931. The repair cassette was amplified using oTO18-oTO19 and was transformed into CauTO496 to generate CauTO550.

**pTO324/CauTO546:** The plasmid backbone was amplified from pTO139 using oTO590 and oTO591. The NAT cassette was amplified from pTO137 using oTO874 and oTO668. The NAT cassette was flanked by 500bp of regions homologous to the regions immediately upstream and downstream of *RPN4* (B9J08_003372) amplified from *C. auris* genomic DNA using oTO1983-oTO1984 and oTO1985-oTO1986, respectively. The repair cassette was amplified using oTO18 and oTO19 and transformed into AR0382 to generate CauTO546.

**pTO346/CauTO604:** The plasmid backbone was amplified from pTO139 using oTO590 and oTO591. The NAT cassette was amplified from pTO137 using oTO874 and oTO668. The NAT cassette was flanked by 500bp of regions homologous to the regions immediately upstream and downstream of *PRE9* (B9J08_002268) amplified from *C. auris* genomic DNA using oTO2125-oTO2126 and oTO2133-oTO2134, respectively. The repair cassette was amplified using oTO18 and oTO19 and transformed into AR0382 to generate CauTO604.

**pTO370/CauTO629:** The plasmid backbone was amplified from pTO139 using oTO590 and oTO591. The ADH1 terminator and NEO cassette was amplified from pTO169 using oTO1066 and oTO668. The NEO cassette was flanked upstream by the ORF of *PRE9* (B9J08_002268) and 500bp of upstream sequence, which was amplified from *C. auris* genomic DNA using oTO2125 and oTO2210. A downstream homologous flanking sequences was amplified from *C. auris* genomic DNA using oTO2133 and oTO2134. The repair cassette was amplified using oTO18 and oTO19 and transformed into CauTO604 to generate CauTO629.

**pTO325/CauTO552:** The entire pTO306 vector was amplified in two overlapping fragments using overlapping backbone-specific primers oTO1427 and oTO1428 paired with oTO1958 and oTO1959, respectively. oTO1958 and oTO1959 contain the C-terminus of POC4 with an ALFA epitope tag directly before the stop codon. The repair cassette was amplified using oTO18-oTO19 and transformed into CauTO467 to generate CauTO552.

**pTO341/CauTO591:** The *POC4* complementation vector was amplified from pTO306 using oTO2106 and oTO1066. A CuG-optimized and C-terminal ALFA-tagged *POC4* gene based off *S. cerevisiae* S288C *POC4* was obtained as a gene fragment from Twist Biosciences (San Francisco, California, USA) and assembled with the amplified fragment to generate pTO341. The repair cassette was amplified using oTO18-oTO19 and transformed into CauTO467 to generate CauTO591.

**pTO350/CauTO611:** 1000bp of promoter sequence upstream of *TEF1* (B9J08_003610) was amplified from *C. auris* genomic DNA using oTO1168 and oTO1169. The entire pTO325 cassette was amplified with pTO2178 and oTO2179 and assembled to incorporate p*TEF1* directly upstream of the *ScPOC4* ORF. The repair cassette was amplified using oTO18 and oTO19 and transformed into CauTO467 to generate CauTO611.

**pTO352/CauTO612:** 1000bp of promoter sequence upstream of *TEF1* (B9J08_003610) was amplified from *C. auris* genomic DNA using oTO1168 and oTO1169. The entire pTO325 cassette was amplified with oTO2179 and oTO2184 and assembled to incorporate p*TEF1* directly upstream of the *POC4* ORF. The repair cassette was amplified using oTO18 and oTO19 and transformed into CauTO467 to generate CauTO612.

**pTO381/CauTO689:** The IRC25 deletion cassette was amplified from pTO315 using oTO1427 and oTO2343 and combined with the NEO cassette from pTO322 amplified using oTO669 and oTO1428. The repair cassette was amplified using oTO18 and oTO19 and transformed into CauTO467 to generate CauTO689.

**pTO435/CauTO804/CauTO805/CauTO806/CauTO807/CauTO808:** The IRC25 complementation cassette was amplified into three fragments to incorporate a C-terminal 4xFLAG construct. The first fragment was amplified from pTO322 using oTO2717 and oTO2749, while the second fragment was amplified using oTO2748 and oTO2718. The HYG resistance cassette was amplified using oTO2719 and oTO2720. The repair cassette was amplified using oTO18 and oTO19 and transformed into CauTO496 to generate CauTO804. The repair cassette was transformed into CauTO552, CauTO611, CauTO799 and CauTO800 to generate CauTO805, CauTO806, CauTO807, and CauTO808, respectively.

**pTO421/CauTO799:** The *POC4* high-expression complementation vector was amplified from pTO350 using oTO1169 and oTO1066. The Seq2523 protein was synthesized as a CUG-optimized gene fragment containing a C-terminal ALFA tag from Twist Biosciences and assembled with the complementation vector to generate pTO421. The repair cassette was amplified using oTO18 and oTO19 and transformed into CauTO467 to generate CauTO799.

**pTO422/CauTO800:** The *POC4* high-expression complementation vector was amplified from pTO350 using oTO1169 and oTO1066. The Seq2212 protein was synthesized as a CUG-optimized gene fragment containing a C-terminal Alfa tag from Twist Biosciences and assembled with the complementation vector to generate pTO422. The repair cassette was amplified using oTO18 and oTO19 and transformed into CauTO467 to generate CauTO800.

**pTO423/CauTO801:** The *POC4* high-expression complementation vector was amplified from pTO350 using oTO1169 and oTO1066. The Seq1379 protein was synthesized as a CUG-optimized gene fragment containing a C-terminal Alfa tag from Twist Biosciences and assembled with the complementation vector to generate pTO423. The repair cassette was amplified using oTO18 and oTO19 and transformed into CauTO467 to generate CauTO801.

**pTO424/CauTO802:** The *POC4* high-expression complementation vector was amplified from pTO350 using oTO1169 and oTO1066. The Seq2350 protein was synthesized as a CUG-optimized gene fragment containing a C-terminal Alfa tag from Twist Biosciences and assembled with the complementation vector to generate pTO424. The repair cassette was amplified using oTO18 and oTO19 and transformed into CauTO467 to generate CauTO802.

**pTO425/CauTO803:** The *POC4* high-expression complementation vector was amplified from pTO350 using oTO1169 and oTO1066. The Seq1823 protein was synthesized as a CUG-optimized gene fragment containing a C-terminal Alfa tag from Twist Biosciences and assembled with the complementation vector to generate pTO425. The repair cassette was amplified using oTO18 and oTO19 and transformed into CauTO467 to generate CauTO803.

### *C. auris* Transformation

*C. auris* transformation was performed using a transient-Cas9 expression system as described previously (*1*). Transformation repair cassettes were amplified from assembled plasmids. The Cas9 expression cassette was amplified from pTO135 with oTO40 and oTO41. Cassettes for the expression of sgRNA targeting specific loci were amplified from pTO136 using overlap extension PCR to change the gRNA sequence. All linear PCR products were purified using a Zymo DNA Clean & Concentrator kit (Zymo Research, D4034) according to the manufacturer’s instructions.

To prepare competent *C. auris* cells, a single colony was inoculated into liquid YPD and incubated overnight at 30 °C with gentle shaking. Cells were harvested by centrifugation and resuspended in TE Buffer and 100 mM lithium acetate. Cells were further incubated at 30 °C for 1 hour with constant shaking before DTT was added to the cells at a final concentration of 25 mM before further incubating at 30 °C for an additional 30 minutes. Cells were harvested by centrifugation at 4 °C before being washed once with ice-cold water and once with ice-cold 1 M Sorbitol. Harvested cells were resuspended in ice-cold 1 M Sorbitol and kept on ice for immediate use or stored at -80 °C for future use.

Electroporation was performed by adding 50 µL of competent cells to a pre-chilled electro-cuvette (2mm-gap) with 500 ng of each of the PCR-amplified Cas9, sgRNA, and repair cassettes. Cells were electroporated using a Bio-Rad MicroPulser Electroporator using the pre-defined *P. pastoris* (PIC) protocol (2.0 kV, 1 pulse). Electroporated cells were immediately recovered in 1M sorbitol and then resuspended in YPD for a 2-hour outgrowth at 30 °C with constant shaking. Outgrowth cells were plated onto selective media. For repair cassettes containing the *NAT* marker, cells were selected on YPD plates containing 200µg/mL nourseothricin with incubation at 30 °C for 2 days. For repair cassettes containing the *NEO* marker, cells were selected on YPD plates containing 1 mg/mL G418 with incubation at 30 °C for 2 days. For repair cassettes containing the *HYG* marker, cells were selected on YPD plates containing 800 µg/mL Hygromycin B with incubation at 30 °C for 2 days.

Transformant colonies were passaged to isolation and correct incorporation of the repair cassette was confirmed for each by colony PCR using Phire Plant Direct PCR Master Mix (F160; Thermo Fisher Scientific) according to the manufacturer’s instructions. Each mutant was confirmed with at least three independent PCR reactions with distinct primer sets specific to the mutation site and compared to parental strains.

### Phylogenetic Analysis

The protein sequences used in the phylogenetic analysis are available in Table S4. Sequence homologs of the indicated genes were identified using the *S. cerevisiae* amino acid sequence as the query using NCBI BLAST (https://blast.ncbi.nlm.nih.gov/Blast.cgi). For *CauPOC4* homologs, the *C. auris POC4* amino acid sequence was used instead. The sequence was individually queried against each of the listed species using the blastp algorithm in the non-redundant protein sequences (nr) database under default parameters. Hits were identified by having reciprocal best matches and an E-value less than 0.0001. Syntenic genes were identified in the listed species using both the Candida Gene Order Browser and the Yeast Gene Order Browser (*2*, *3*).

For each gene, the homologous sequences were first aligned using muscle5 and trimmed using trimAI (parameters: -gt 0.7 -cons 60) and then manually adjusted to remove unalignable sites from downstream analyses (*4*, *5*). The curated sequence files used in downstream analyses are provided in Table S4. These alignment files were used to infer the maximum likelihood phylogeny for each gene using the best-fit model of sequence evolution and rate variation parameters using the IQTree2 software and the implemented ModelFinder function (*6*, *7*). This method provided a direct estimate of the evolutionary distance between pairs of sequences in the alignment.

### Homology analysis

For sequence-based homology, 5,417 protein coding sequences from FungiDB (version 68) CaurisB8441 were searched using BlastP (Protein-Protein BLAST 2.12.0+) against all 572,970 Swiss-Prot (version 2025_01). For structure-based homology, the same protein coding sequences were mapped to structures in the AFDB (v4, model creation date: 2022-06-01) as follows. First, the sequences were mapped to locus tags in the NCIB GCA_002759435.3_Cand_auris_B8441_V3 genome yielding 5,395 one-to-one matches. Note FungiDB CaurisB8441 ids have the form B9J08_###### (6 numbers), while GCA_002759435.3_Cand_auris_B8441_V3 ids have the form B9J08_##### (5 numbers). Then GCA_002759435.3_Cand_auris_B8441_V3 locus tags were mapped UniPort entries for the UP000230249 Proteome, where all ids mapped 1-1. Each UniProt entry was then mapped to UniParc ids, which uniquely identifies the sequence. Then, the AFDB was queried for structures for *C. auris*, yielding 12,017 structures and their pLDDT scores were recorded. For each AFDB structure the corresponding UniProt entry was mapped its UniParc entry and de-duplicated. Joining *C. auris* protein and AFDB structures by UniParc ids yielded 4,360 AFDB structures for *C. auris* proteins. We then used Foldseek (downloaded 2/2025, Foldseek version 6cfb8805be28925a81f194e4b616ad5616f2ed9a and MMseq version 6cfb8805be28925a81f194e4b616ad5616f2ed9a) to search for structure homologues in the pre-defined Alphafold/Swiss-Prot reference database, using foldseek easy-search command with default parameters otherwise. To compare sequence vs structure homology, for each *C. auris* gene, we computed the maximum BlastP and Foldseek bit score across Swiss-Prot targets.

### Structural Predictions

Protein models of mid assembly proteasome assembly complex were predicted using AlphaFold 3 (https://alphafoldserver.com/ 5/2024) with default parameters. Detailed summary statistics of each model are listed in Table S10. For each structure, the top ranking model was visualized with ChimeraX and Predicted aligned error (PAE) plots were made with PAE Viewer (*8*, *9*). DockQ2 scores were calculated between all chains in a given structure using *ipSAE* as an average across 5 models (*10*). Poc4 chaperone topology was determined using Pro-origami under default parameters (*11*).

### Protein Design

As an initial structure for Poc4, the *C. auris* Poc4 and Irc25 sequences were jointly folded with AlphFold3 (https://alphafoldserver.com/ 5/2024) with default parameters and the top ranking structured was used, having scores pTM: 0.89, ipTM: 0.91, has_clash: 0.0, fraction_disordered: 0.01, and ranking_socre: 0.91. Then given this structure, Frame2Seq (version 0.0.5) was used to generate designs for the Poc4 chain, generated excluding cysteine, varying temperature in [0.01, 0.1, 1, 10], and either using no additional constraints, fixing conserved residues determined by the multiple sequence alignment [18, 24, 26, 42, 44, 52, 57, 67, 76, 78, 79, 83, 85, 86, 89], or alternatively fixing the Poc4 residues at the Poc4-Irc25 interface [1, 3, 5, 12, 14, 16, 18, 24, 25, 26, 28, 30, 32, 33, 34, 35, 36, 37, 38, 39, 42, 44, 55, 56, 57, 58, 83, 85, 87]. From the temperature=1.0 and fixinterface parameters, 5000 sequences were generated, and each was folded with AlphaFold2. ParaFold was used to distribute first the MSA generation and then the folding tasks onto the University of Michigan Great Lakes CPU and GPU HPC cluster queues (*12*). Designs were evaluated by the Frame2Seq Score, AlphaFold2 pLDDT, sequence identity to the native Poc4, and diversity among the designs. To visualize the similarity between designs, Poc4 and structural homologues, ESM2 embeddings for the Poc4 Foldseek hits, members of pfam PF10448, and the designs were generated using the esm2_t33_650M_UR50D model, by averaging the per-residue representation in the 33^rd^ layer for the up to 1022 first residues in the sequence. The EMS2 embeddings were then projected to 2-dimensions using UMAP (v0.5.7) with parameters a=1, b=1.8, metric=”cosine”.

### Growth Assays

Spot assays were performed by growing fungal cells overnight in liquid YPD media at 30 °C with constant shaking. Cell culture density was normalized to an OD_600_ of 1 in sterile water and diluted 10-fold. 5µL of each dilution were spotted onto YPD plates and incubated at 30 °C, 37 °C, or 42 °C for 2 days before imaging on a BioRad gel dock. Each plate image is representative of at least 3 biological replicates.

Kinetic growth assays were performed by growing fungal cells overnight in 200µL of YPD in a 96 well plate at 30 °C, then subcultured into fresh media using a sterile pinner. Growth curves were performed on a BioTek LogPhase600 Plate Reader at the indicated temperatures for 24 hours with shaking at 500 rpm and OD_600_ measurements taken every 10 minutes. A minimum of 4 biological replicates, each with 3 technical replicates was performed per strain per growth condition. Growth curves were plotted in GraphPad Prism v10.

### Minimum Inhibitory Concentration (MIC) Assays

Standard compound sensitivity testing was evaluated by broth microdilution MIC testing in 96 well, flat-bottom microtiter plates. All compounds were dissolved in DMSO. Tested MG-132 (Cayman Chemical, 10012628) and bortezomib (Cayman Chemical, 10008822) concentrations ranged from 3.125 µM to 200µM. Tested fluconazole (Cayman Chemical, 11594), caspofungin (GoldBio, C-520-10), and amphotericin B (GoldBio, A-560-250), concentrations ranged from 0.04-64 µg/mL, 0.08-80 ng/mL, and 0.04-64 µg/mL, respectively. Assays were set up in a total volume of 200 µL of YPD per well with 2-fold serial dilutions of the test compound. Following inoculation, plates were covered and incubated statically at the indicated temperature. Growth was quantified by measuring OD_600_ with a BioTek 800 TS Absorbance Reader at an endpoint of 24 hours. Relative growth was calculated for each strain’s by dividing the optical density at a given concentration, by the optical density of an untreated control on the same plate. All strains were assessed with 3 biological replicates in technical duplicates. Dose-response curves were plotted in GraphPad Prism v10.

### Co-Immunoprecipitation

Strains were inoculated in YPD and subcultured to an OD_600_ of 0.05 in 100 mL of fresh YPD. Cultures were grown for 6 hours at 37 °C in 250 mL Erlenmeyer flasks with 150 rpm shaking until mid-log phase. Cells were collected by centrifugation at 3500rpm, washed in 1mL sterile water, transferred to 1.5mL screw-cap tubes, then flash frozen and stored at -80 °C.

Cell pellets were resuspended in 1mL of lysis buffer (20 mM Tris pH 7.5, 100 mM KCl, 5 mM MgCl, 20% glycerol, and 1X HALT protease inhibitor cocktail (Thermo Fisher Scientific, 87786) added fresh before use). The tube was filled with acid-washed glass beads until the beads were just below the meniscus at the top of the tube to limit foaming during bead beating. Cells were disrupted by bead beating three times for 4 minutes, with 7-minute incubation on ice between cycles. Lysates were recovered by a stacked transfer and clarified by centrifugation at 14,000 rpm for 10 min at 4 °C. Protein concentrations were determined by Bradford assay.

Anti-ALFA immunoprecipitations were performed with ALFA Selector PE magnetic agarose beads (NanoTag Biotechnologies, N1515). For each reaction, 20 µL of magnetic agarose bead slurry was washed 2 times in 1 mL ice-cold lysis buffer. Beads were collected by magnetic separation between each wash step. 1mL of protein lysates normalized to a total protein level of 2mg and added to the washed beads. Reactions were incubated with end-over-end rotation for 2 hours at 4 °C. The samples were then washed 3 times in 1mL ice-cold lysis buffer, transferred to clean tubes, and washed again twice with 1 mL ice-cold lysis buffer, and one final time with 1mL ice-cold Tris-buffered saline (TBS). The washed beads were then resuspended in 100 µL of Laemmli sample buffer, incubated at 95 °C for 5 min, shaking at 500 rpm. Supernatant was collected and subject to western immunoblotting on SDS-PAGE gels.

### Protein Extraction

Strains were inoculated overnight in YPD and subcultured to an OD_600_ of 0.1 in 10 mL of fresh media, incubated for 6 hours with shaking at 30 °C. Following sub-culture, cells were harvested by centrifugation, then washed once with sterile water. Cells were resuspended in lysis buffer (50 mM HEPES pH 7.4, 150 mM NaCl, 5 mM EDTA, 1% Triton X-100, 100mM NaF, 1 mM PMSF, 1X HALT protease inhibitor cocktail (Thermo Scientific, 87786)) and lysed by bead beating. Following lysis, lysates were clarified by centrifugation and protein concentration was determined by Bradford assay.

### Western Blotting

Protein levels were normalized to 25 µg and boiled in 1x Laemmli sample buffer for 5 minutes before loading on 4-20% SDS-PAGE gels (Bio-Rad, #4561094). Following separation, proteins were transferred to polyvinylidene difluoride (PVDF) membrane (Bio-Rad, #162-0177) for 15 minutes using a semi-dry transfer system. Blots were blocked with 5% bovine serum albumin (Sigma-Aldrich, A7030) in phosphate-buffered saline with 0.1% Tween-20 (PBS-T). The molecular weight was approximated by electrophoresis of marker proteins in either the PageRuler pre-stained protein ladder (Thermo Scientific, 26616) or PageRuler Plus pre-stained protein ladder (Thermo Scientific, 26619). ALFA epitopes were detected using camelid anti-ALFA Licor 800CW primary antibody (NanoTag Biotechnologies, N1502-Li800-L) at a 1:5000 dilution in blocking solution. FLAG epitopes were detected using 1:1000 dilution rabbit anti-FLAG primary antibody (Sigma-Aldrich, F7425-.2MG). Blots were washed with PBS-T and incubated with 1:15000 goat anti-rabbit IgG IRDye-680RD (LI-COR Biosciences, 926-68071) or 1:15000 goat anti-rabbit IgG IRDye-800CW (LI-COR Biosciences, 926-32211). Ubiquitin epitopes were detected with 1:1000 dilution mouse anti-Ubiquitin primary antibody (Invitrogen, 13-1600). Blots were washed with PBS-T and incubated with 1:15000 dilution anti-mouse IgG IRDye-800CW secondary antibody (LI-COR Biosciences, 926-33210) before detecting signals using a LI-COR Odyssey scanner. Protein levels were normalized to total protein loading using Revert 700 total protein stain (LI-COR Biosciences, 926-11011). Proteasome α-subunits were detected using a primary antibody specific to a conserved peptide in the 20S alpha proteasome proteins Pre1, Pre2, Pre3, Pre5, Pre6, and Pre7 at a 1:3000 dilution in blocking solution (AbCam, antibody ID: AB_2171376). Following washing, blots were incubated with 1:15000 dilution goat anti-mouse IgG HRP secondary antibody (EDM Millipore, 12-348) before detecting signals using a ThermoFisher iBright 1500 imager.

### Statistics and Reproducibility

Statistical analyses were carried out using GraphPad Prism v10. Data are presented as mean ± standard deviation from biological replicates. Unless otherwise noted, experiments were performed in at least three independent biological replicates yielding similar results. Statistical significance among different groups was calculated using one-way ANOVA, with Dunnett’s post hoc tests for multiple comparisons. *p ≤ 0.05; **p ≤ 0.01; ***p ≤ 0.001; n.s., p > 0.05.

---

**Fig. S1.**
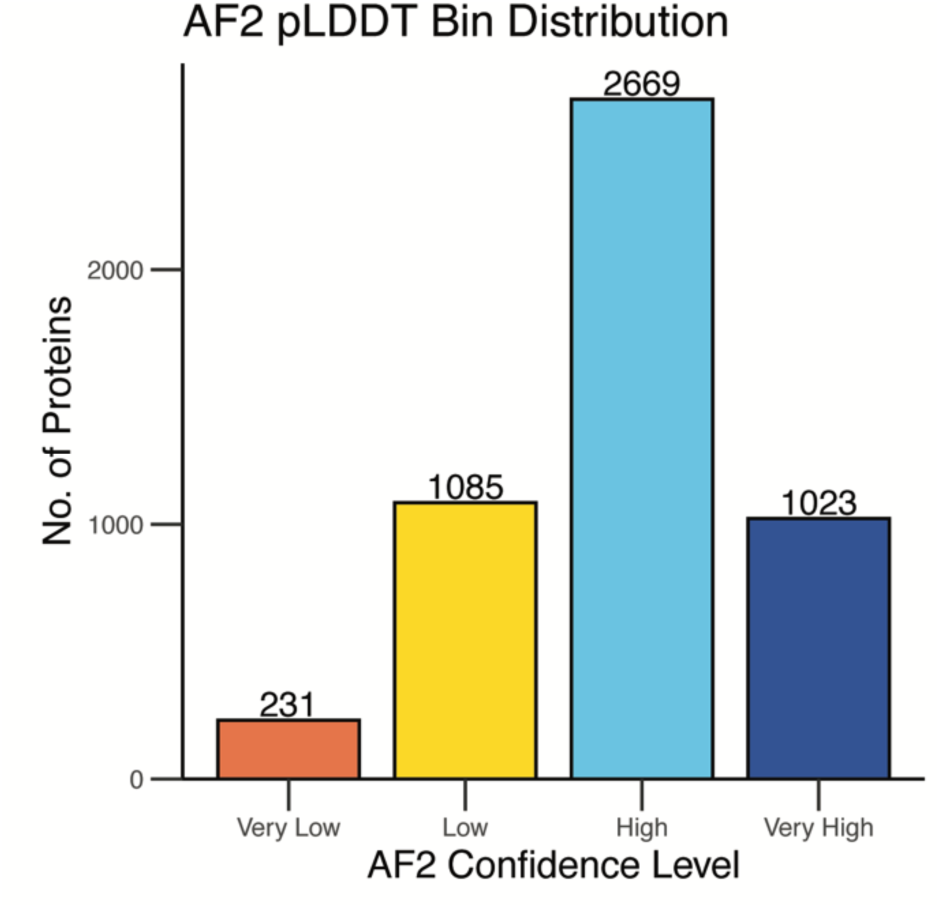
Histogram of global pLDDT of available *C. auris* AlphaFold2 predicted structures. Number above bar indicates number of structures in a given category. Bins are as follows: Very Low < 50; Low 50-70; High 70-90; Very High > 90. Structures were obtained from AlphaFold2 Protein Structure Database (*13*).

**Fig. S2.**
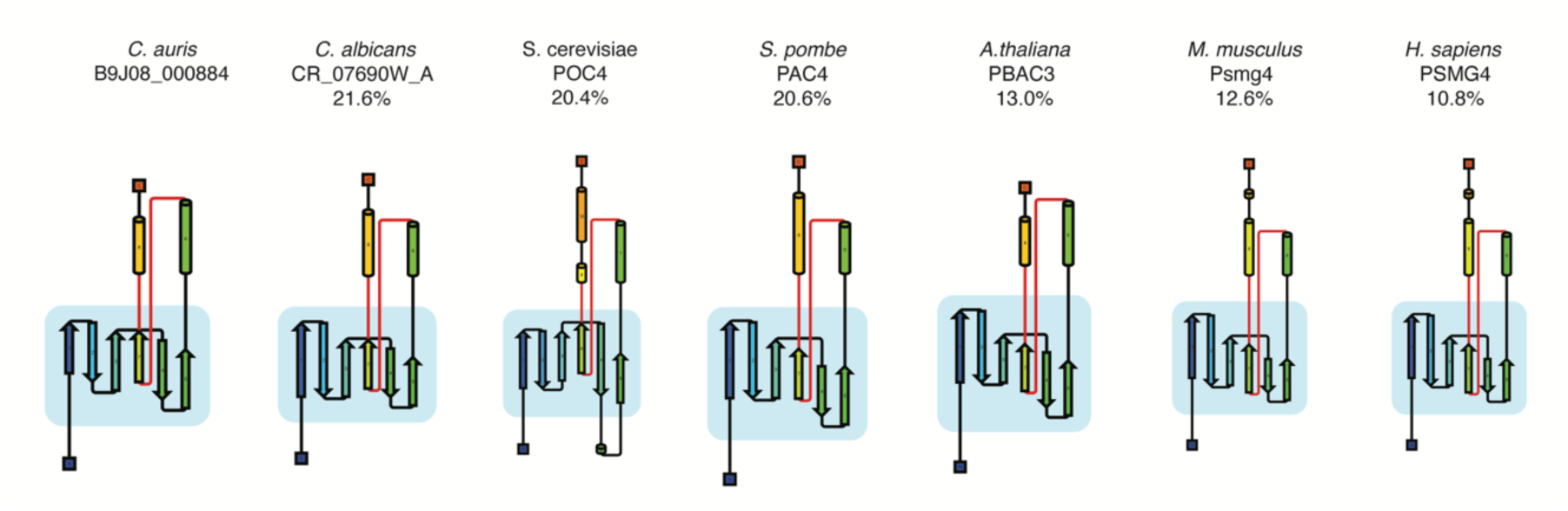
Topology of protein structures from Figure 1F. Arrows indicate sheets, while cylinders represent helices. Percentage indicates amino acid identity to *C. auris* B9J08_000884.

**Fig. S3.**
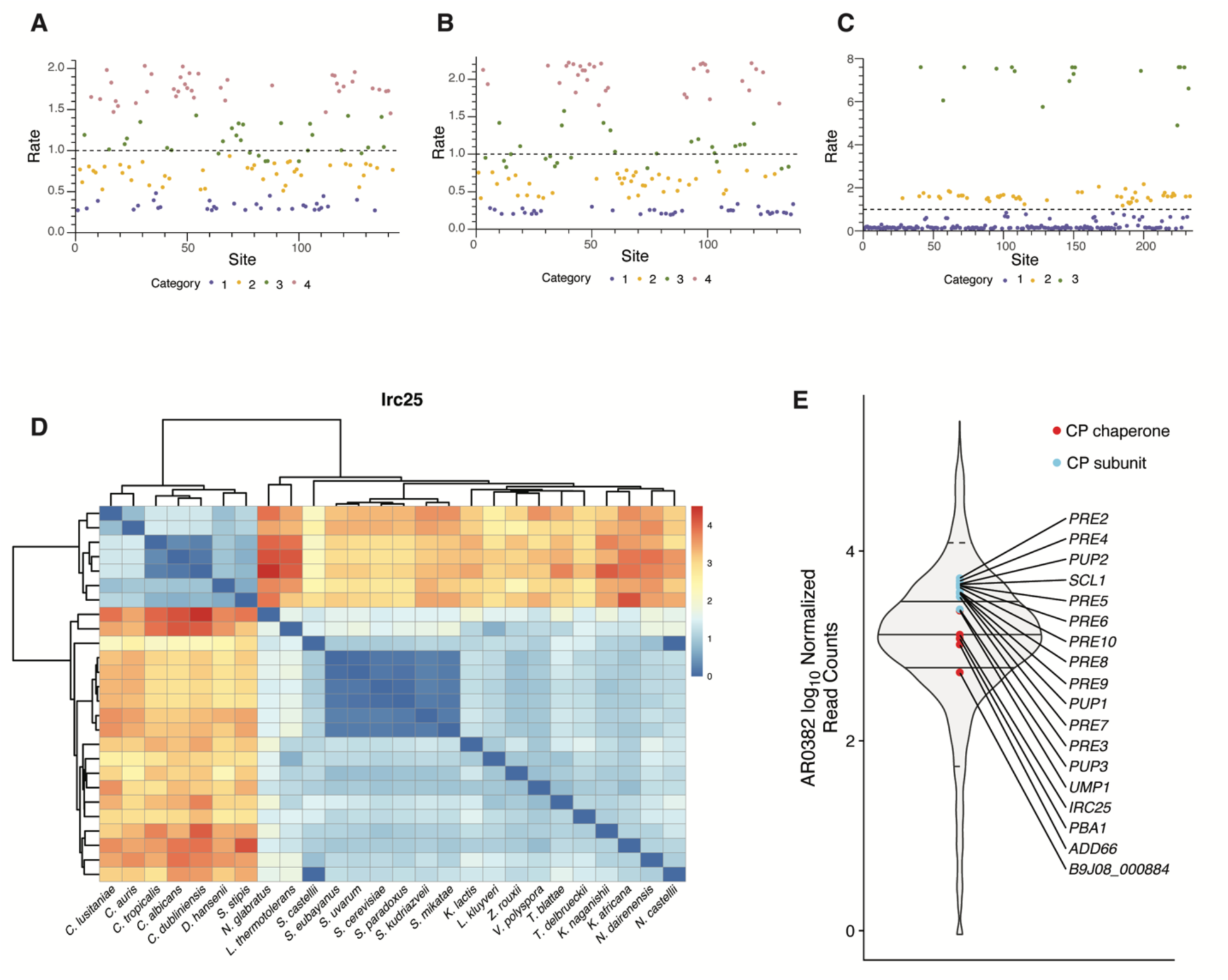
Evolutionary rate variation by residue for **(A)** Poc4, **(B)** Irc25, and **(C)** Pup2. **(D)** Pairwise comparisons of evolutionary rate between identified Irc25 sequence orthologs from indicated species. **(E)** DESeq-normalized read counts of all AR0382 ORFs obtained from (*14*). Solid lines represent IQR, while dashed lines represents 5 and 95 percentiles.

**Fig. S4.**
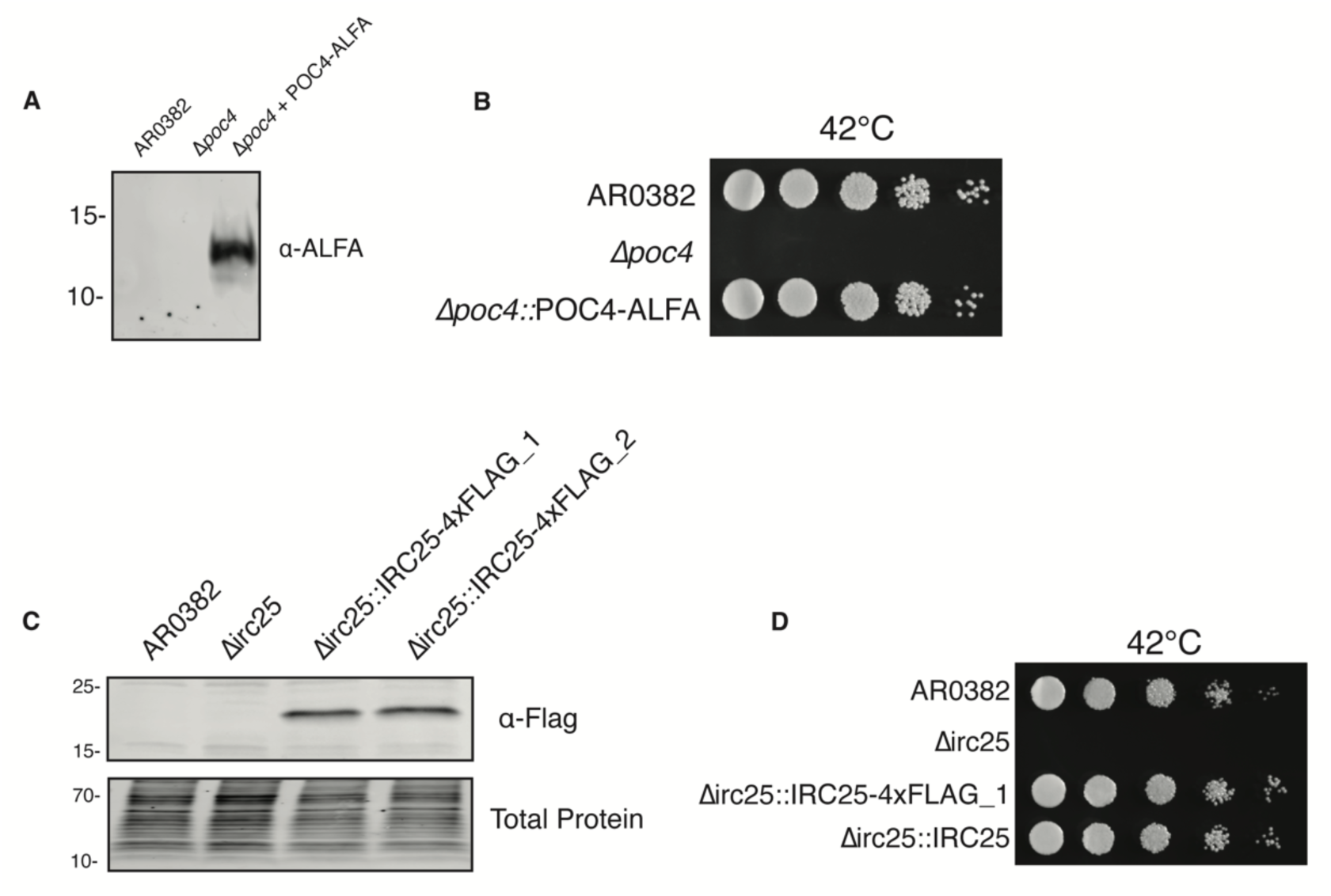
Validation that Poc4-Alfa and Irc25-4xFlag constructs are functional. Western blots showing specific band at expected size in tagged strains for **(A)** Poc4 and **(C)** Irc25. Total protein is shown as a loading control. Spot plates showing full rescue of thermotolerance upon complementation with tagged complements for **(B)** Poc4 and **(D)** Irc25. For spot plates, 10-fold dilutes were incubated on YPD at the indicated temperature for 48 hours before imaging. All images are representative of three biological replicates.

**Fig. S5.**
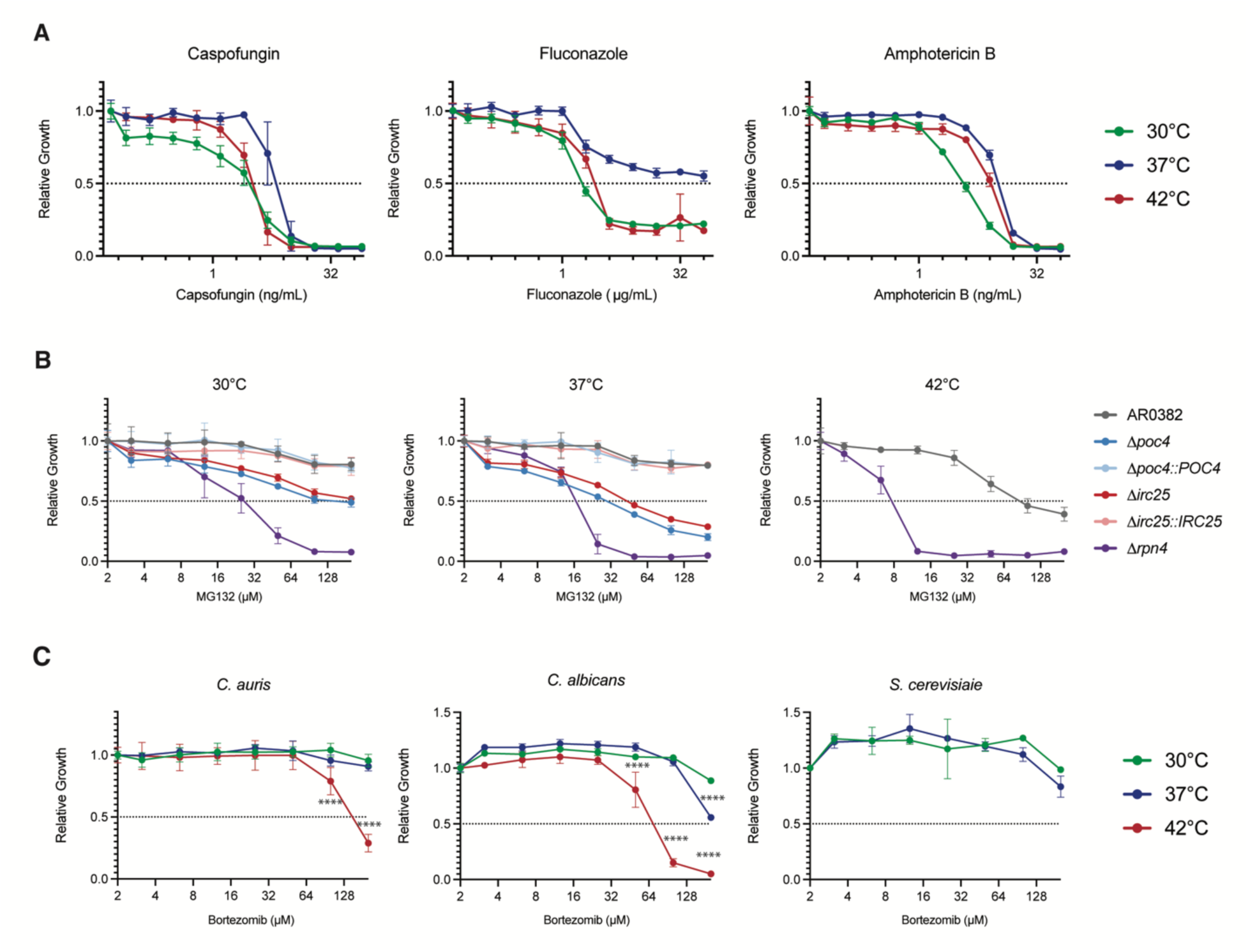
**(A)** Dose-response curves of *C. auris* to the antifungals caspofungin, fluconazole, and amphotericin B at the indicated temperatures. **(B)** Dose-response curves of indicated *C. auris* assembly chaperone mutants show hypersensitivity to MG-132 at all tested temperatures. **(C)** Dose-response curves of *C. auris*, *C. albicans*, and *S. cerevisiae* to Bortezomib at the indicated temperatures. Growth measured by OD_600_ at each drug concentration is normalized to the growth of each strain in the absence of drug to determine relative growth. Data show mean ± standard deviation across 3 biological replicates. Statistical significance among different groups was calculated using one-way ANOVA, with Dunnett’s post hoc tests for multiple comparisons. *p ≤ 0.05; **p ≤ 0.01; ***p ≤ 0.001; n.s., p > 0.05.

**Fig. S6.**
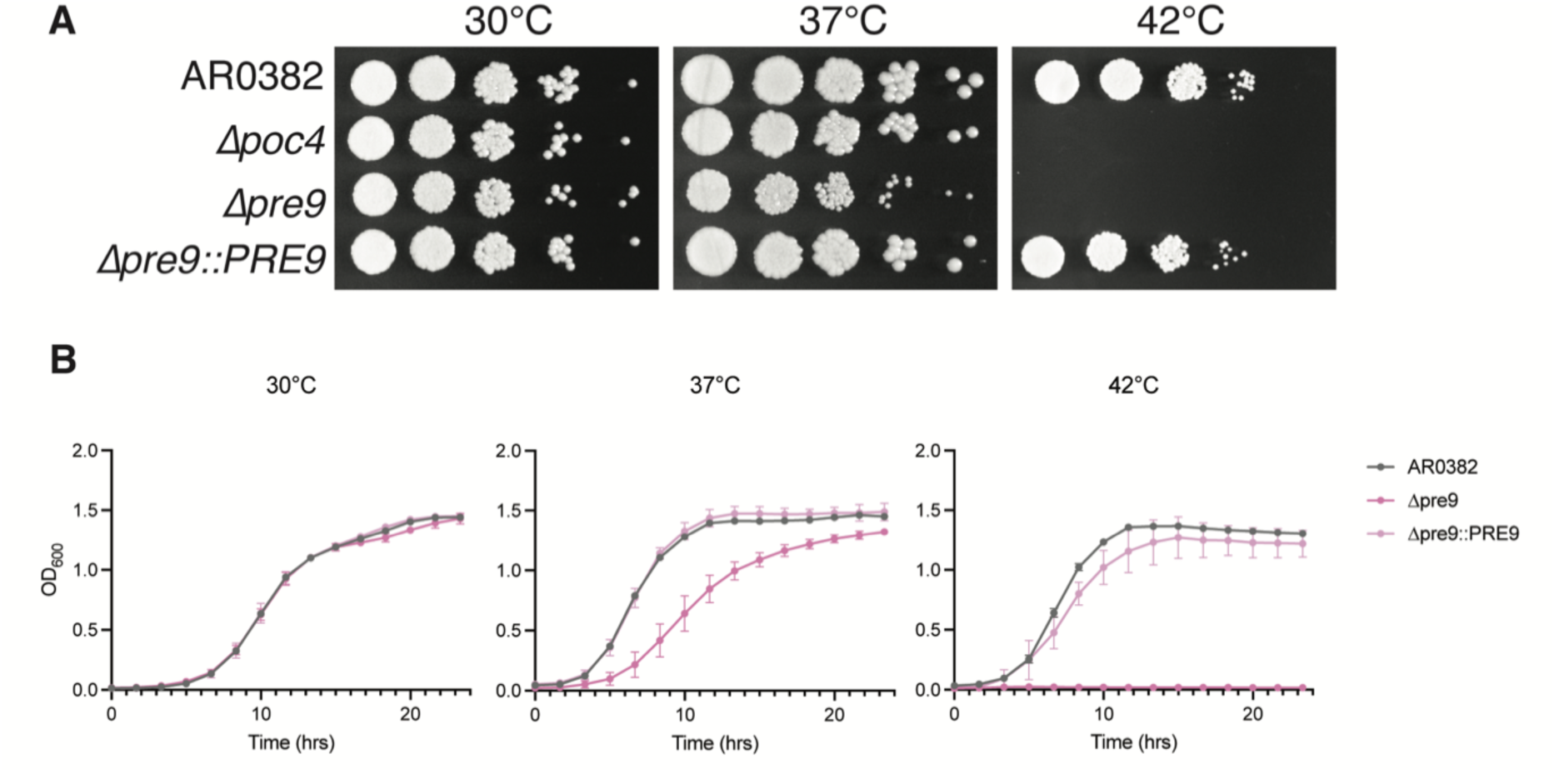
Loss of proteasome subunit Pre9 results in more extreme temperature sensitivity than loss of Poc4 or Irc25. **(A**) Spot plates of indicated *C. auris* strains grown at 30 °C, 37 °C, and 42 °C. 10-fold dilutions were incubated on YPD at the respective temperature for 48 hours before imaging. **(B)** Growth curves of indicated *C. auris* strains at 30 °C, 37 °C, and 42 °C. Growth was measured by OD_600_ every 10 minutes for 24 hours. Data shown is the mean ± standard deviation across 4 biological replicates every 80 minutes for clarity. Images are representative of three biological replicates.

**Fig. S7.**
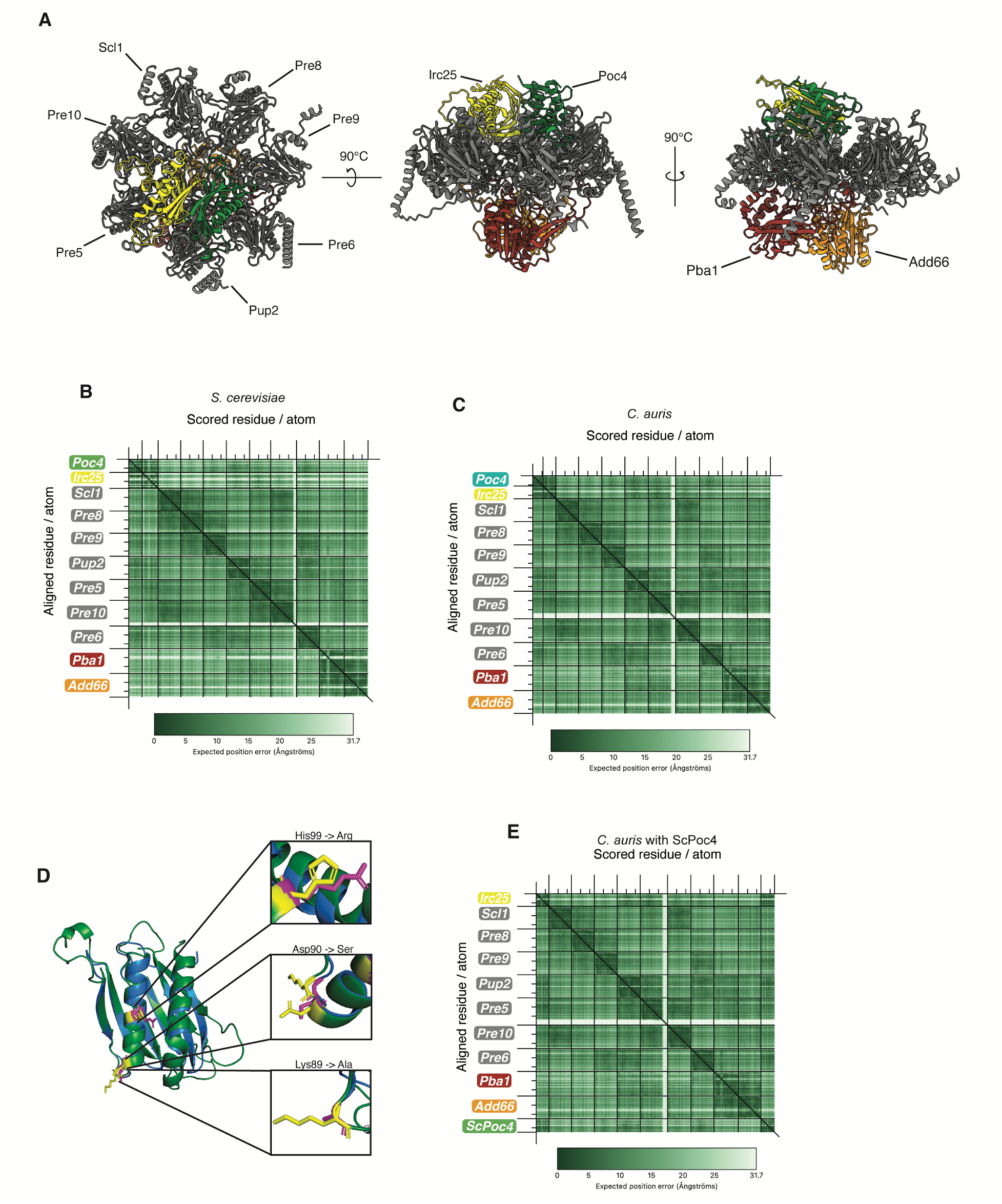
**(A)** AlphaFold 3 model of *S. cerevisiae* proteasome subunits during core particle assembly. Predicted aligned error (PAE) plots, which show the predicted position error at residue x if the predicted and true structures were aligned on residue y, for the AlphaFold 3 models depicted in **(B)** Fig. 4A, **(C)** Fig. **(D)** S7A, and Fig. 4C. Large ticks demarcate individual chains, with minor ticks every 100 residues. **(E)** Aligned predicted structures of *C. auris* Poc4 (blue) and *S. cerevisiae* Poc4 (green). Residues required for binding with Pup2 in *S. cerevisiae* are colored pink and shown in insets. Corresponding residues in *C. auris* are colored yellow.

**Fig. S8.**
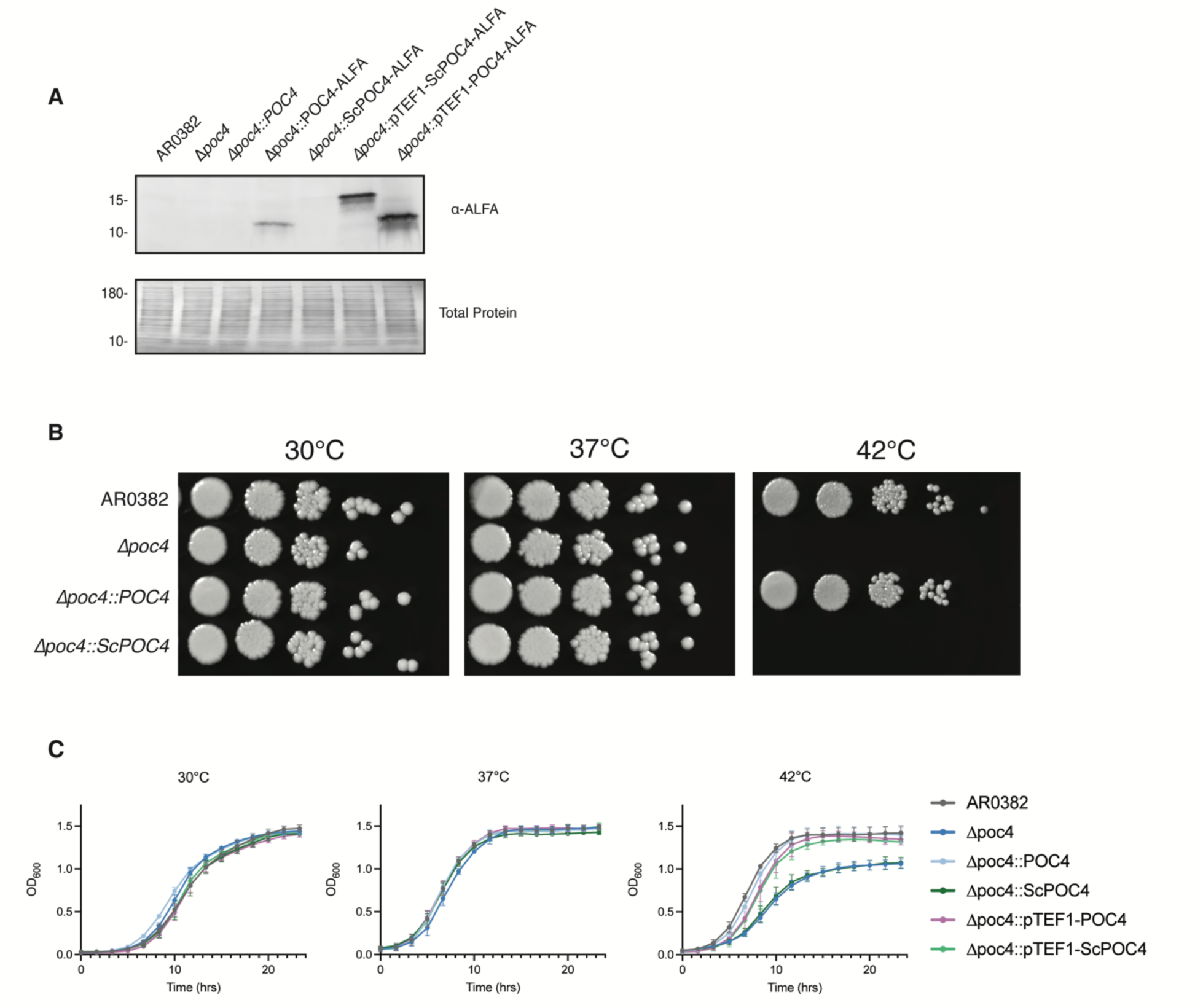
**(A)** Western blot showing expression of various Poc4 constructs in *C. auris.* Indicated strains were grown in fresh media for 5 hours at 30 °C before protein extraction and western blotting. Total protein is shown as a loading control. **(B)** Spot plates of indicated *C. auris* strains grown at 30°C, 37°C, and 42 °C. 10-fold dilutions were incubated on YPD at respective temperature for 48 hours before imaging. **(C)** Growth curves of indicated *C. auris* strains at 30°C, 37°C, and 42 °C. Growth was measured by OD_600_ every 10 minutes for 24 hours. Data shown is the mean ± standard deviation across 4 biological replicates every 80 minutes for clarity. Images are representative of three biological replicates.

**Fig. S9.**
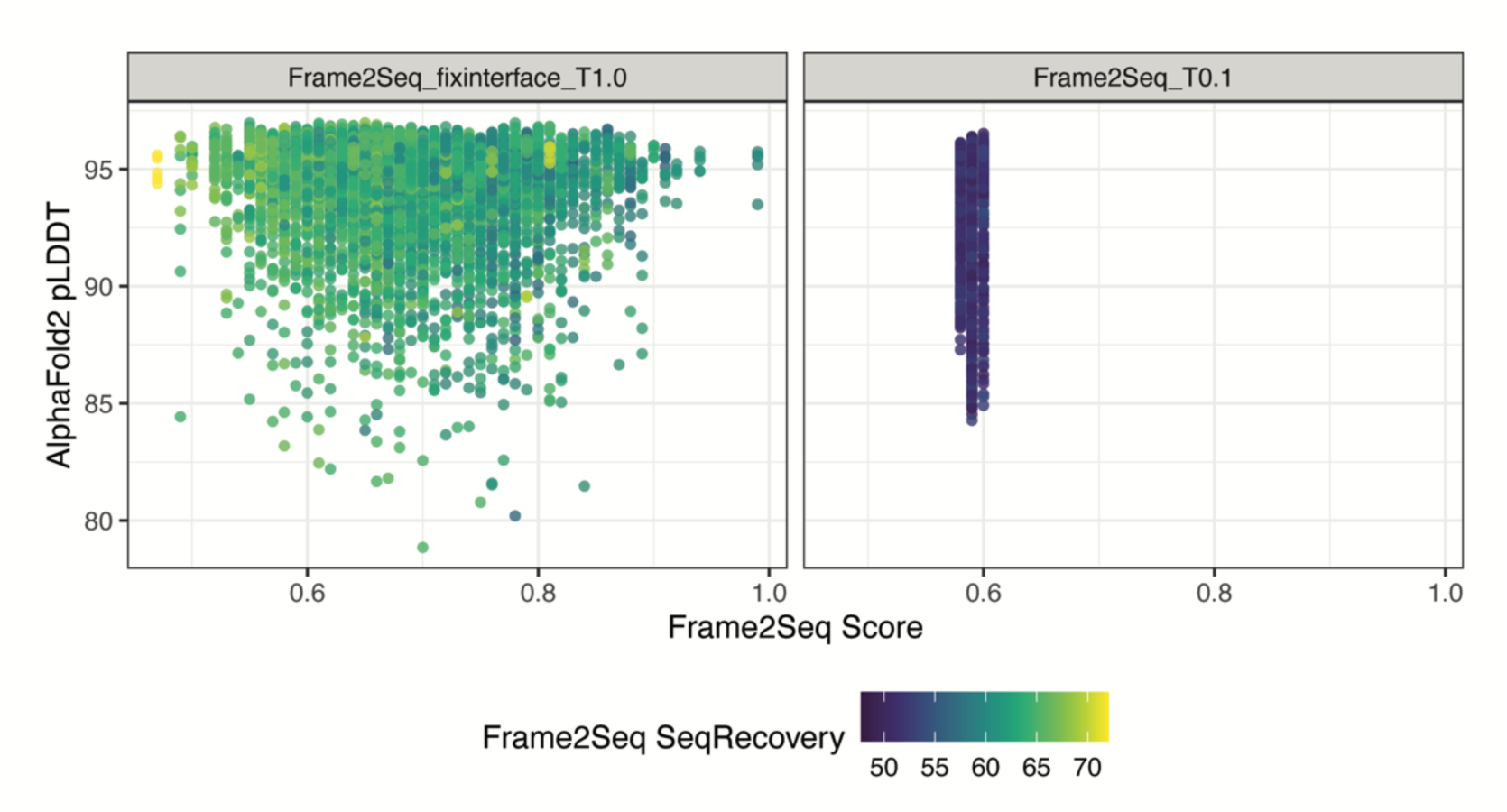
Given the AlphaFold3 predicted structure of the Poc4-Irc25 dimer, we fixed the Irc25 sequence and 29 residues of Poc4 at the interface and excluded the cysteine residue type. We used Frame2Seq with a temperature factor of 1 to design the remaining positions of Poc4 to generate 5,000 designs. Each design was then re-folded with AlphaFold2. Shown is the distribution of the AlphaFold2 pLDDT scores (higher is better) by the Frame2Seq score (lower is better) colored by the percent identity over the designed positions. A substantial fraction of the predicted structures have a high pLDDT score (>90) and sequence recovery ranging between ∼50 and ∼70 percent identity. The Frame2Seq score has low correlation with sequence recovery and no correlation with pLDDT. Together this suggests that the Poc4 scaffold is highly designable by Frame2Seq.

**Fig. S10.**
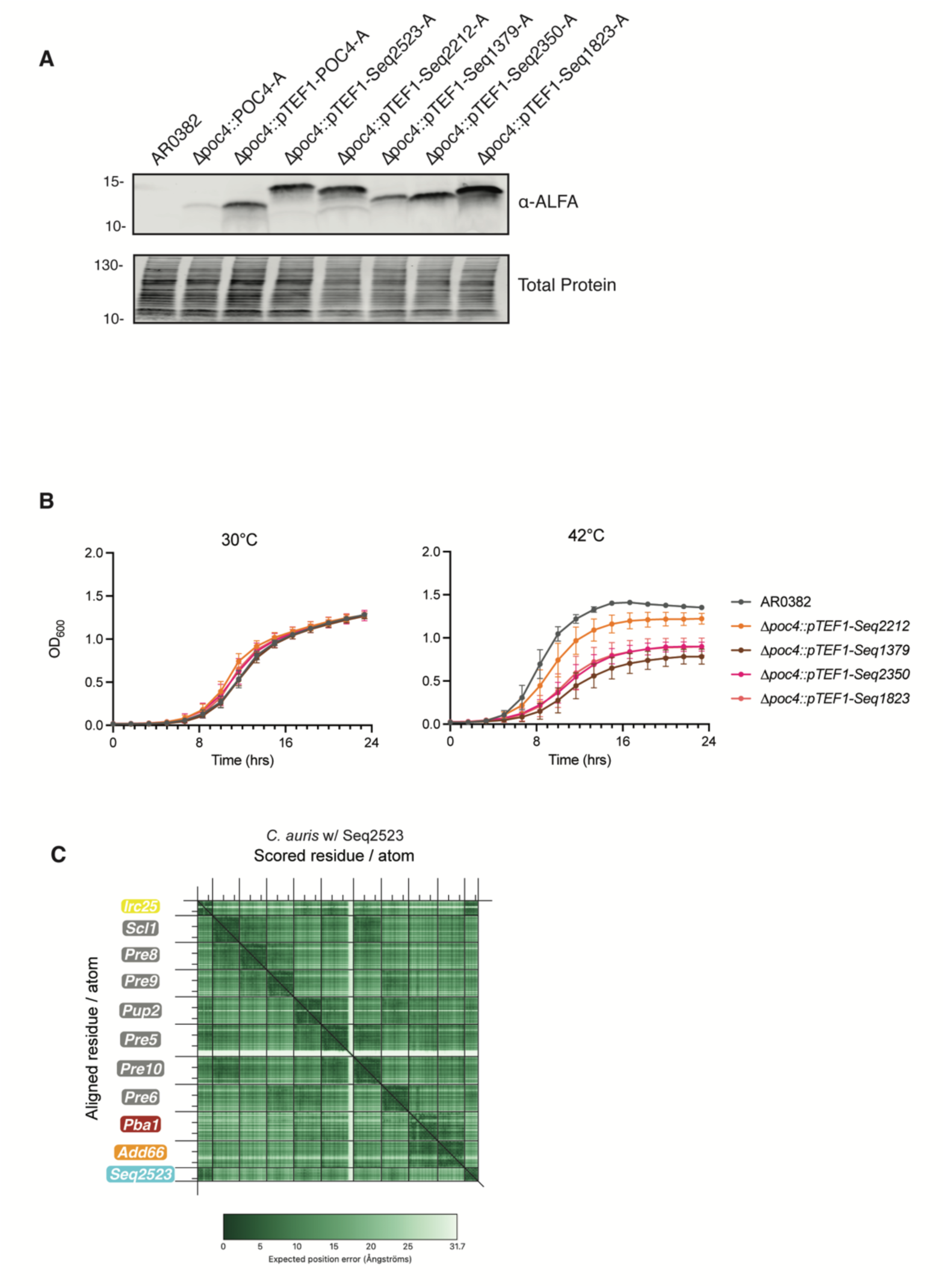
**(A)** Western blot showing expression of indicated chaperone designs in *C. auris*. Strains were grown in fresh media for 5 hours at 30 °C before protein extraction and western blotting. Total protein is shown as a loading control. Image is representative of three biological replicates **(B)** Growth curves of indicated *C. auris* strains at 30 °C, 37 °C and 42 °C. Growth was measured by OD_600_ every 10 minutes for 24 hours. Data shown is the mean ± standard deviation across 4 biological replicates every 80 minutes for clarity. **(C)** PAE plot of AlphaFold 3 model of structure depicted in Fig. 5D.

**Table S1.** List of strains used in this study.

| Strain Name | Background | Genotype | Source |
| --- | --- | --- | --- |
| CauTO33 | *C. auris* B11109 | AR0382 | (15) |
| CaTO1 | *C. ablicans* SC5314 | SC5314 | (16) |
| ScTO499 | *S. cerevisiae* BY4743 | BY4743 | (17) |
| CauTO467 | *C. auris* B11109 | AR0382<br>$\Delta poc4(B9J08\ 000884)::NAT$ | This Study |
| CauTO497 | *C. auris* B11109 | AR0382 $\Delta poc4::NAT::POC4\ NEO$ | This Study |
| CauTO496 | *C. auris* B11109 | AR0382<br>$\Delta irc25(B9J08\ 003917)::NAT$ | This Study |
| CauTO550 | *C. auris* B11109 | AR0382 $\Delta irc25::NAT::IRC25\ NEO$ | This Study |
| CauTO546 | *C. auris* B11109 | AR0382<br>$\Delta rpn4(B9J08\ 003372)::NAT$ | This Study |
| CauTO552 | *C. auris* B11109 | AR0382 $\Delta poc4::NAT::POC4\text{-}ALFA\ NEO$ | This Study |
| CauTO604 | *C. auris* B11109 | AR0382<br>$\Delta pre9(B9J08\ 002268)::NAT$ | This Study |
| CauTO629 | *C. auris* B11109 | AR0382 $\Delta pre9::NAT::PRE9\ NEO$ | This Study |
| CauTO689 | *C. auris* B11109 | AR0382 $\Delta poc4::NAT\ \Delta irc25::NEO$ | This Study |
| CauTO591 | *C. auris* B11109 | AR0382 $\Delta poc4::NAT::ScPOC4\text{-}ALFA\ NEO$ | This Study |
| CauTO611 | *C. auris* B11109 | AR0382 $\Delta poc4::NAT::pTEF1\text{-}ScPOC4\text{-}ALFA\ NEO$ | This Study |
| CauTO612 | *C. auris* B11109 | AR0382 $\Delta poc4::NAT::pTEF1\text{-}CauPOC4\text{-}ALFA\ NEO$ | This Study |
| CauTO804 | *C. auris* B11109 | AR0382 $\Delta irc25::NAT::IRC25\text{-}4xFLAG\ HYG$ | This Study |
| CauTO805 | *C. auris* B11109 | AR0382 $\Delta poc4::NAT::POC4\text{-}ALFA\ NEO\ IRC25::IRC25\text{-}4xFLAG\ HYG$ | This Study |
| CauTO806 | *C. auris* B11109 | AR0382 $\Delta poc4::NAT::pTEF1\text{-}ScPOC4\text{-}ALFA\ NEO\ IRC25::IRC25\text{-}4xFLAG\ HYG$ | This Study |
| CauTO799 | *C. auris* B11109 | AR0382 $\Delta poc4::NAT::pTEF1\text{-}Seq2523\text{-}ALFA\ NEO$ | This Study |
| CauTO800 | *C. auris* B11109 | AR0382 $\Delta poc4::NAT::pTEF1\text{-}Seq2212\text{-}ALFA\ NEO$ | This Study |
| CauTO801 | *C. auris* B11109 | AR0382 $\Delta poc4::NAT::pTEF1\text{-}Seq1379\text{-}ALFA\ NEO$ | This Study |
| CauTO802 | *C. auris* B11109 | AR0382 $\Delta poc4::NAT::pTEF1\text{-}Seq2350\text{-}ALFA\ NEO$ | This Study |
| CauTO803 | *C. auris* B11109 | AR0382 $\Delta poc4::NAT::pTEF1\text{-}Seq1823\text{-}ALFA\ NEO$ | This Study |
| CauTO807 | *C. auris* B11109 | AR0382 $\Delta poc4::NAT::pTEF1$-<br>Seq2523-ALFA NEO<br>IRC25::IRC25-4xFLAG HYG | This Study |
| CauTO808 | *C. auris* B11109 | AR0382 $\Delta poc4::NAT::pTEF1$-<br>Seq2212-ALFA NEO<br>IRC25::IRC25-4xFLAG HYG | This Study |

**Table S2.**
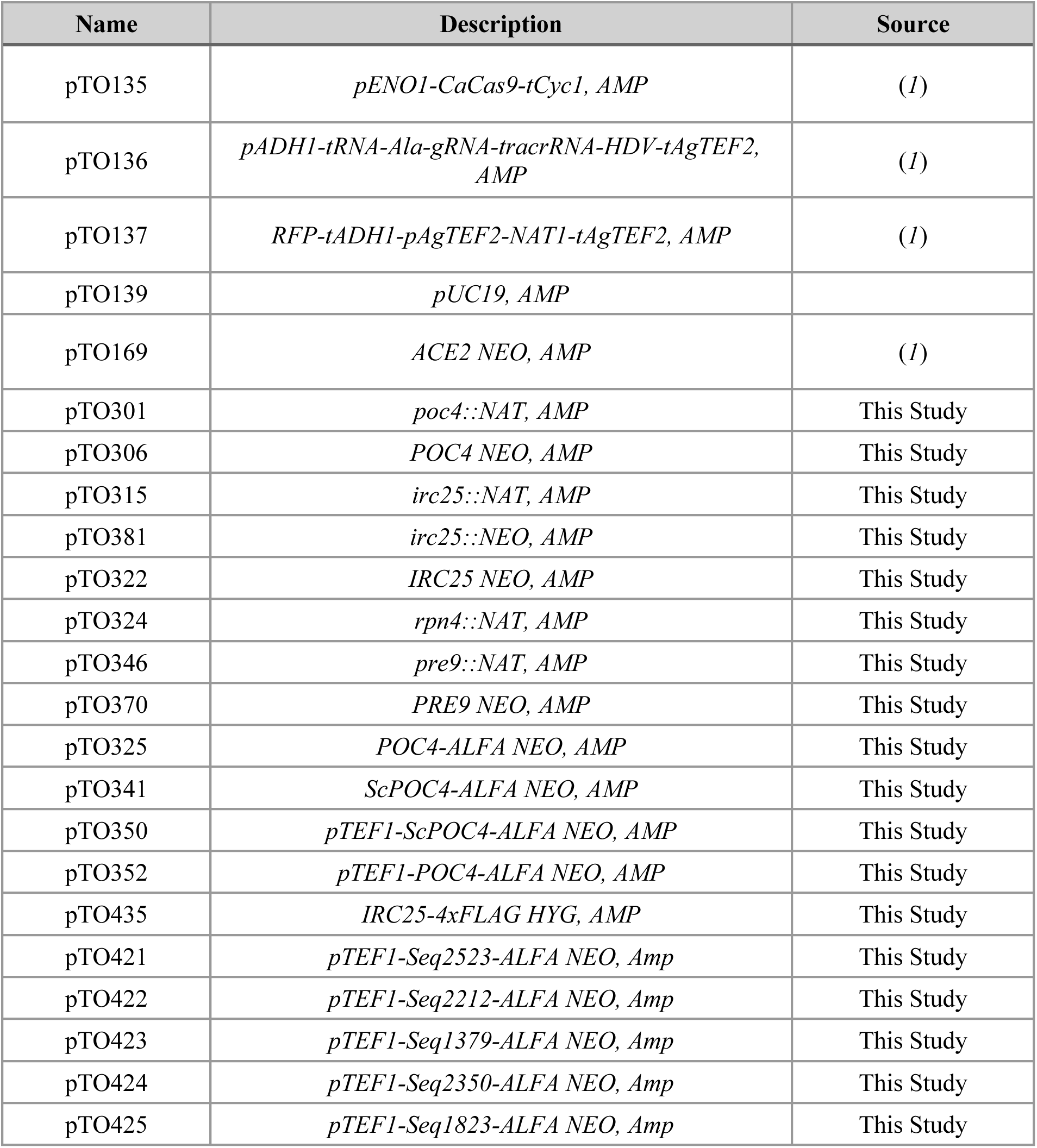
List of plasmids used in this study.

| Name | Description | Source |
| --- | --- | --- |
| pTO135 | *pENO1-CaCas9-tCyc1, AMP* | (1) |
| pTO136 | *pADH1-tRNA-Ala-gRNA-tracrRNA-HDV-tAgTEF2, AMP* | (1) |
| pTO137 | *RFP-tADH1-pAgTEF2-NAT1-tAgTEF2, AMP* | (1) |
| pTO139 | *pUC19, AMP* |  |
| pTO169 | *ACE2 NEO, AMP* | (1) |
| pTO301 | *poc4::NAT, AMP* | This Study |
| pTO306 | *POC4 NEO, AMP* | This Study |
| pTO315 | *irc25::NAT, AMP* | This Study |
| pTO381 | *irc25::NEO, AMP* | This Study |
| pTO322 | *IRC25 NEO, AMP* | This Study |
| pTO324 | *rpn4::NAT, AMP* | This Study |
| pTO346 | *pre9::NAT, AMP* | This Study |
| pTO370 | *PRE9 NEO, AMP* | This Study |
| pTO325 | *POC4-ALFA NEO, AMP* | This Study |
| pTO341 | *ScPOC4-ALFA NEO, AMP* | This Study |
| pTO350 | *pTEF1-ScPOC4-ALFA NEO, AMP* | This Study |
| pTO352 | *pTEF1-POC4-ALFA NEO, AMP* | This Study |
| pTO435 | *IRC25-4xFLAG HYG, AMP* | This Study |
| pTO421 | *pTEF1-Seq2523-ALFA NEO, Amp* | This Study |
| pTO422 | *pTEF1-Seq2212-ALFA NEO, Amp* | This Study |
| pTO423 | *pTEF1-Seq1379-ALFA NEO, Amp* | This Study |
| pTO424 | *pTEF1-Seq2350-ALFA NEO, Amp* | This Study |
| pTO425 | *pTEF1-Seq1823-ALFA NEO, Amp* | This Study |

**Table S3.**
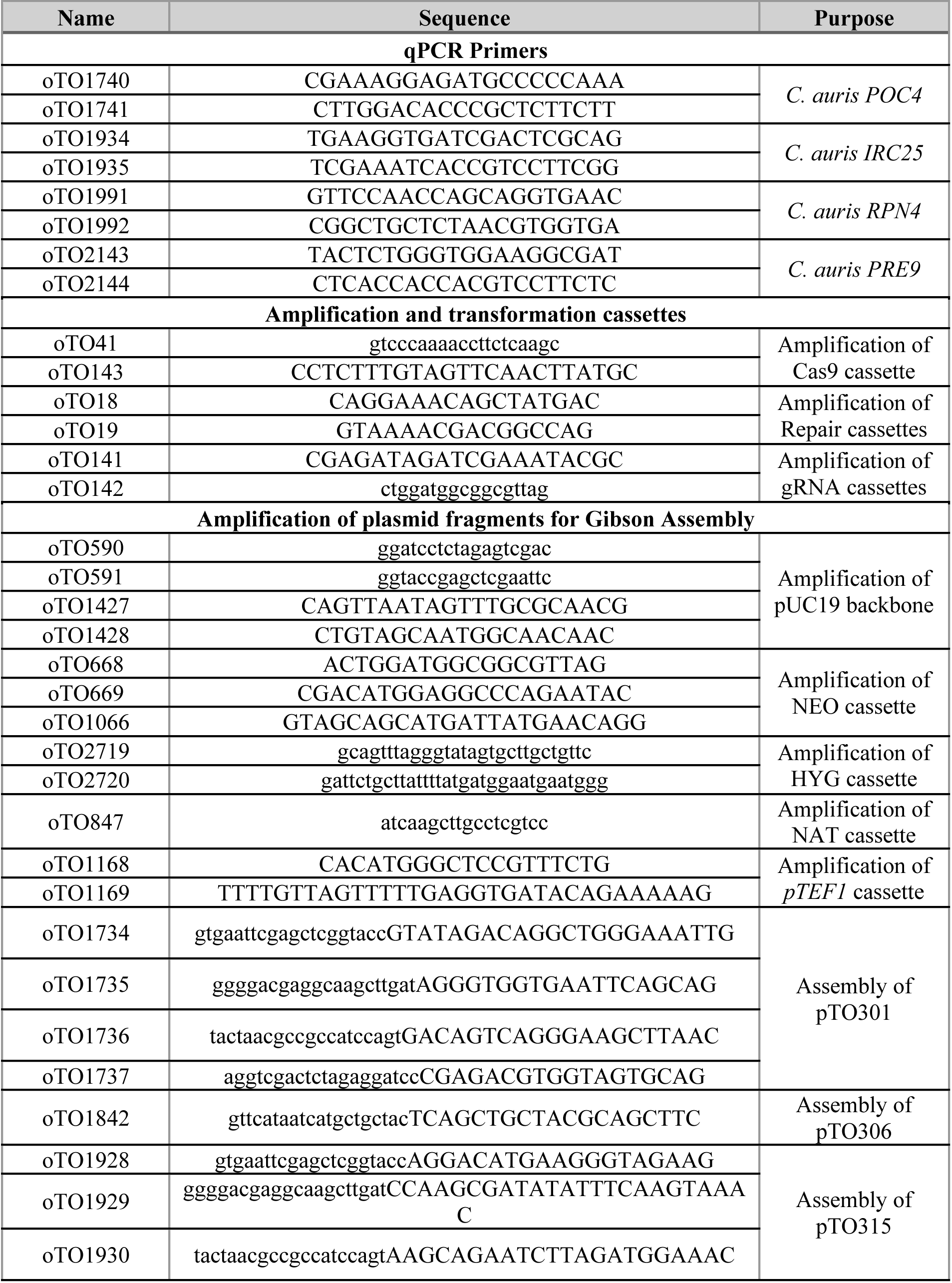
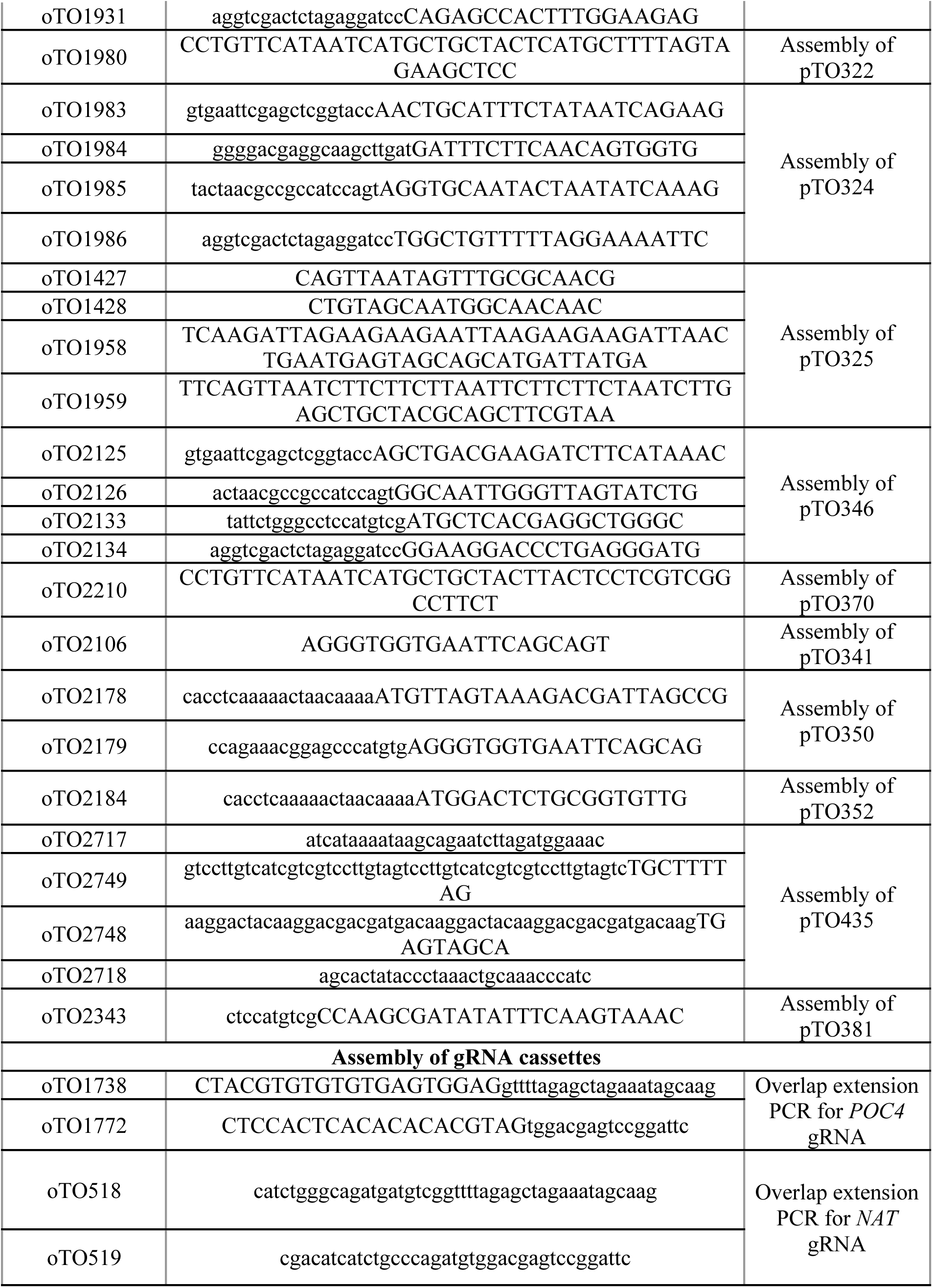
List of oligos used in this study.

**Table S4.**
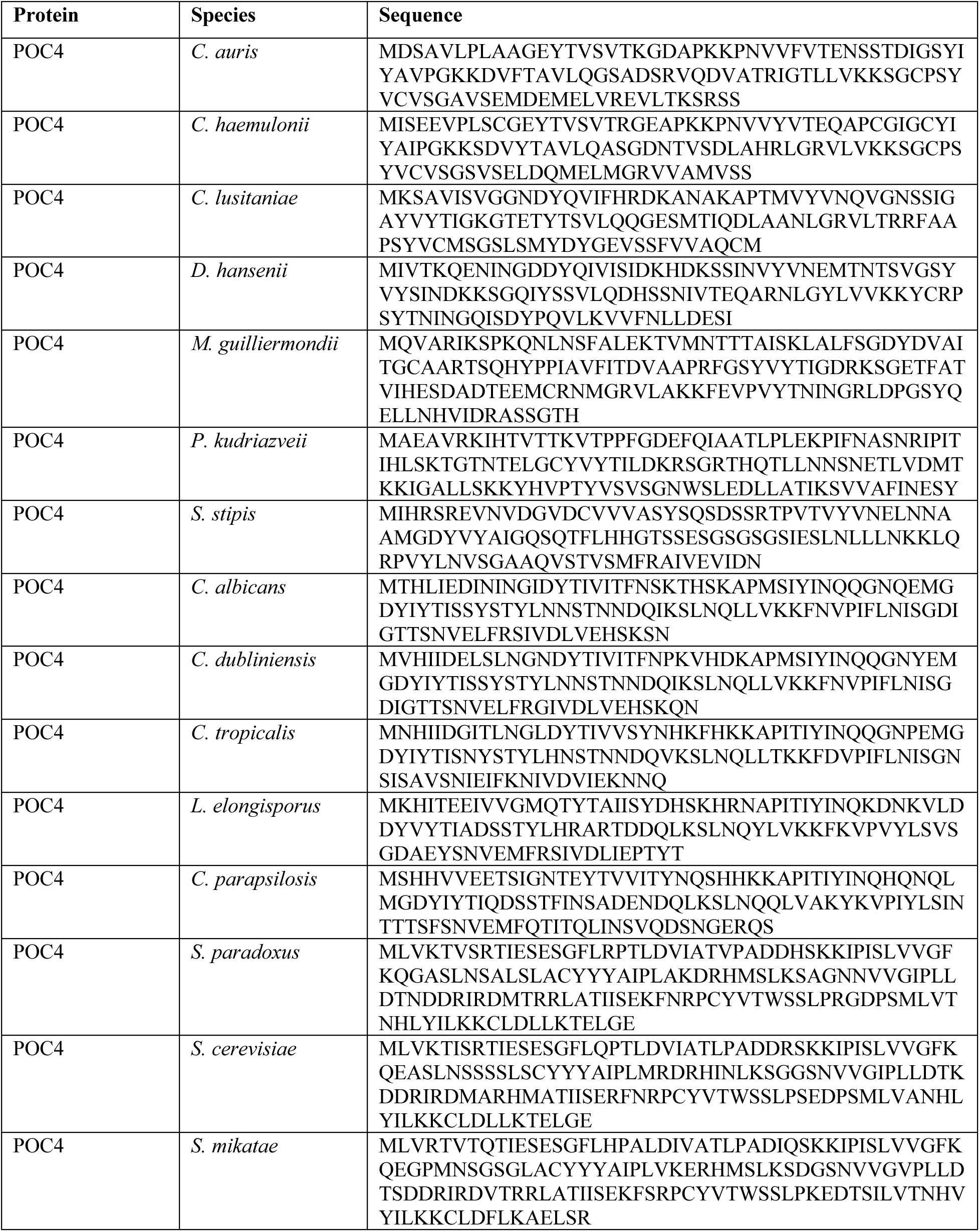
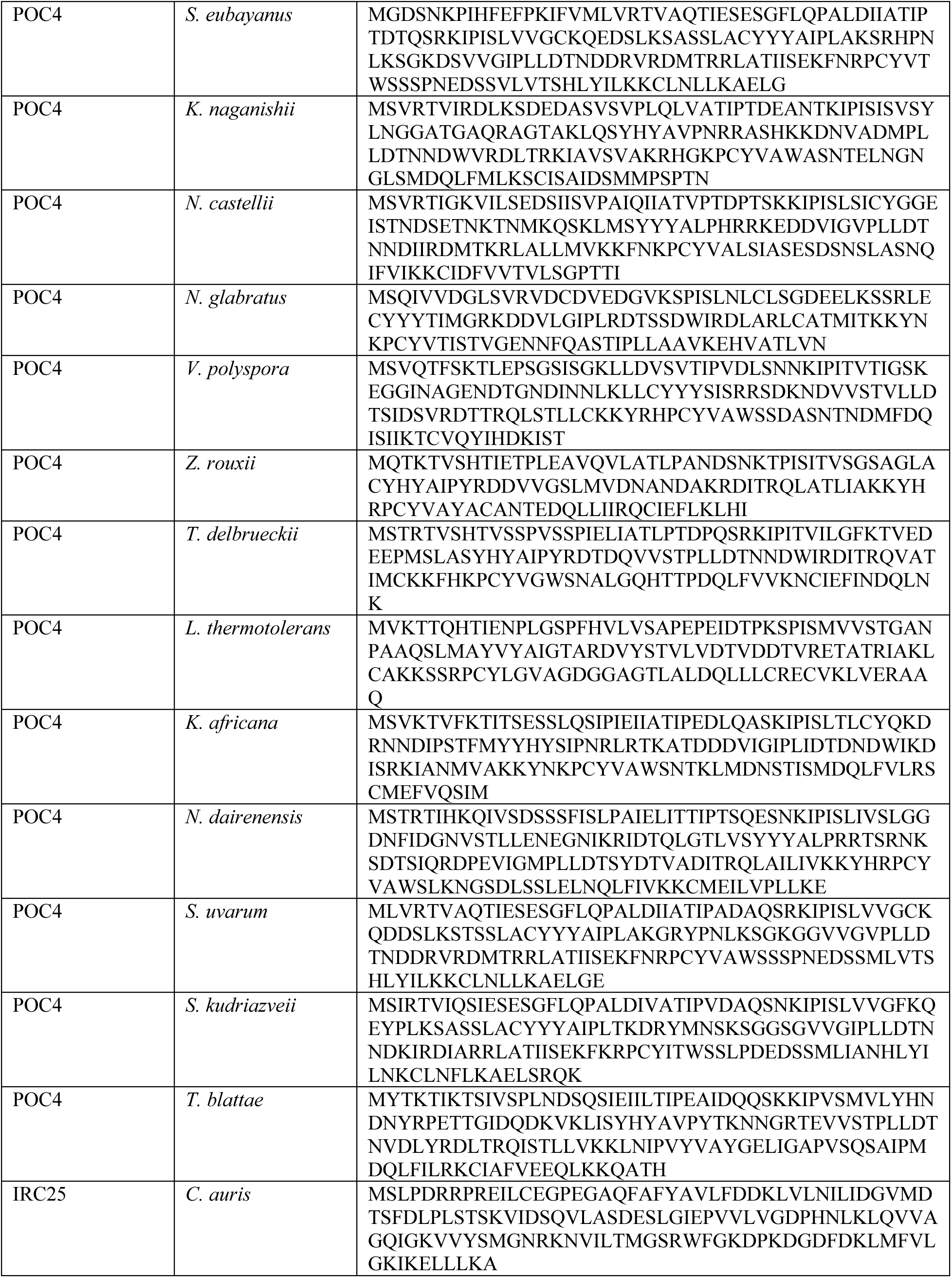
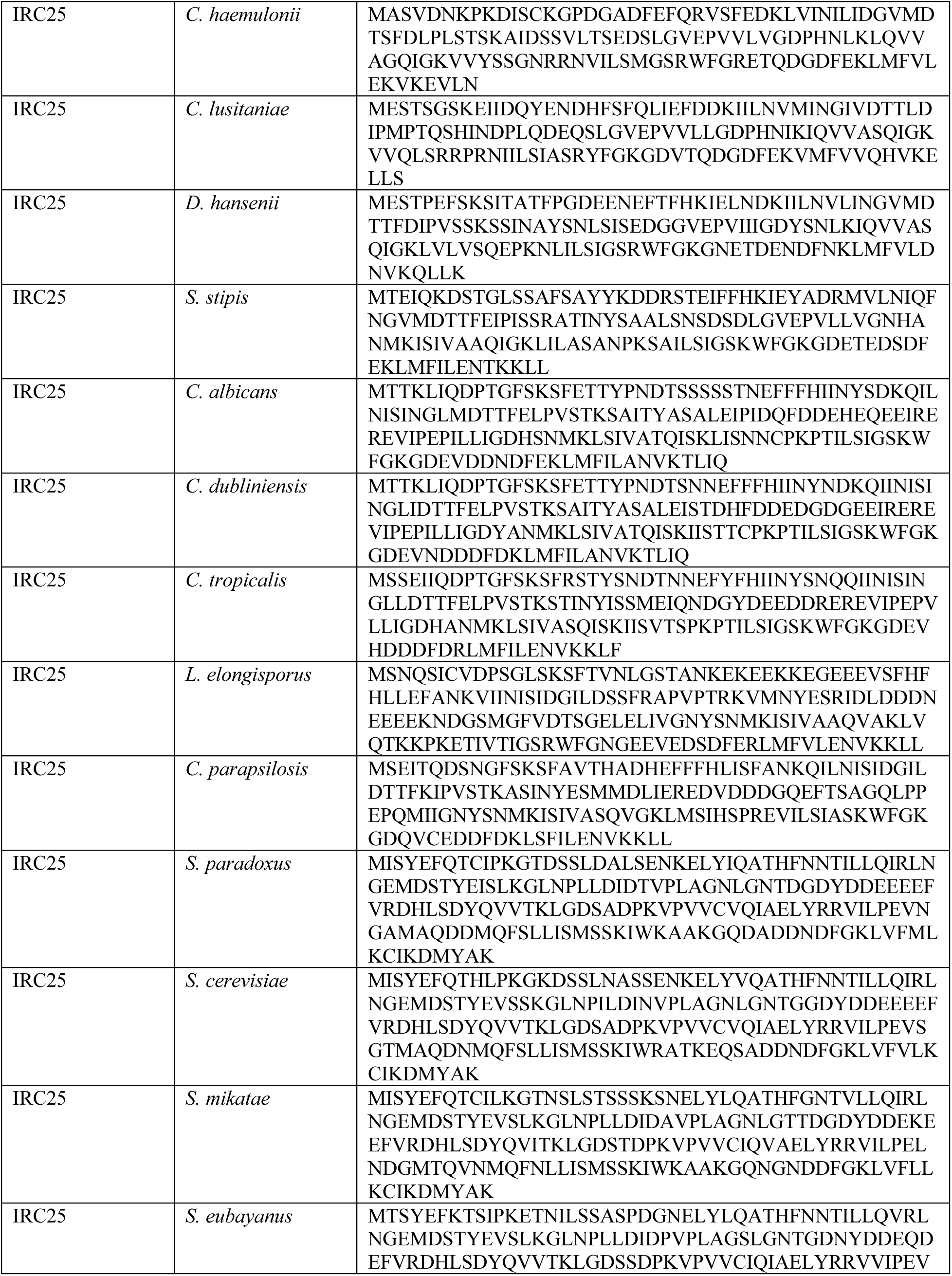
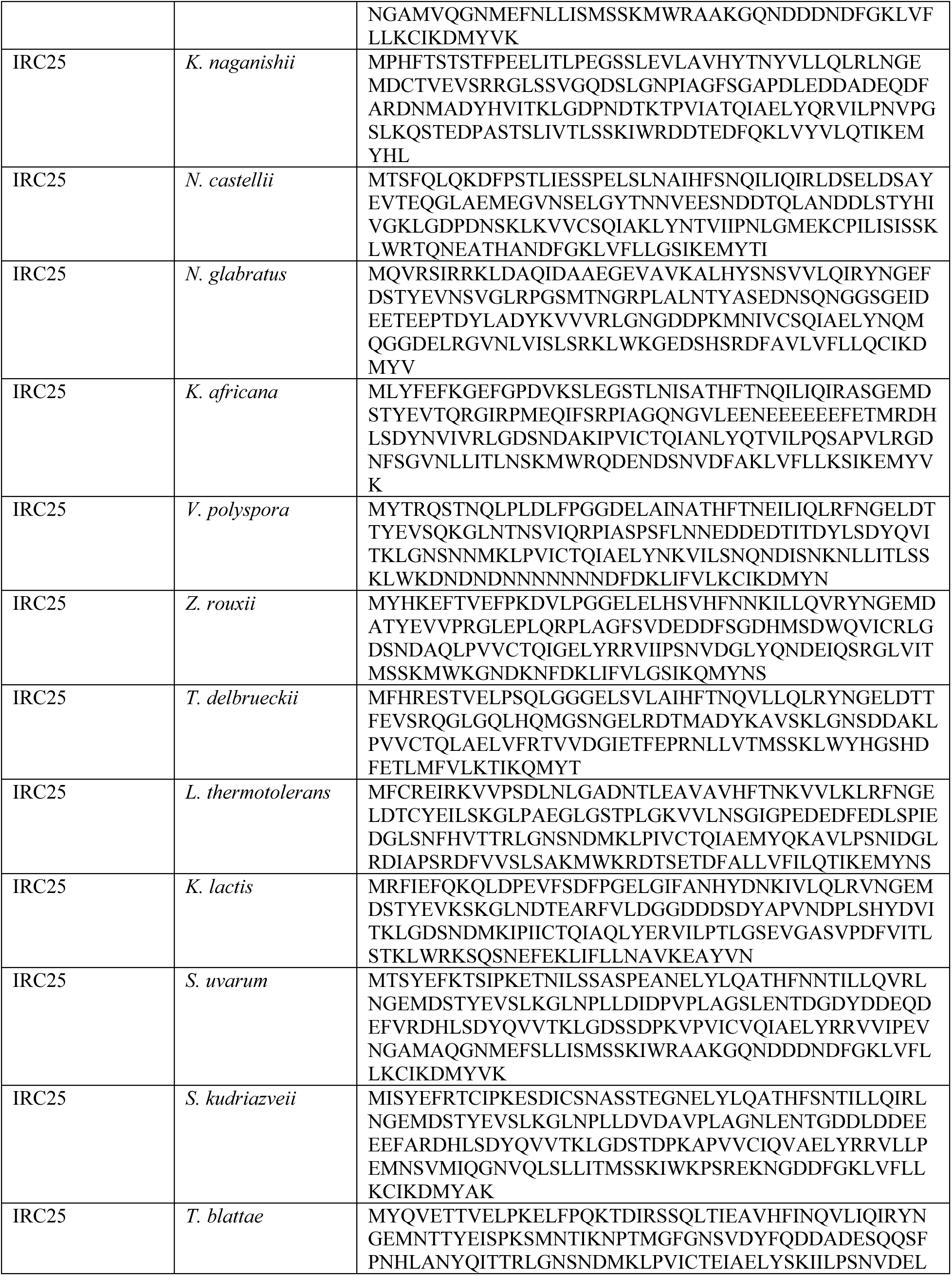
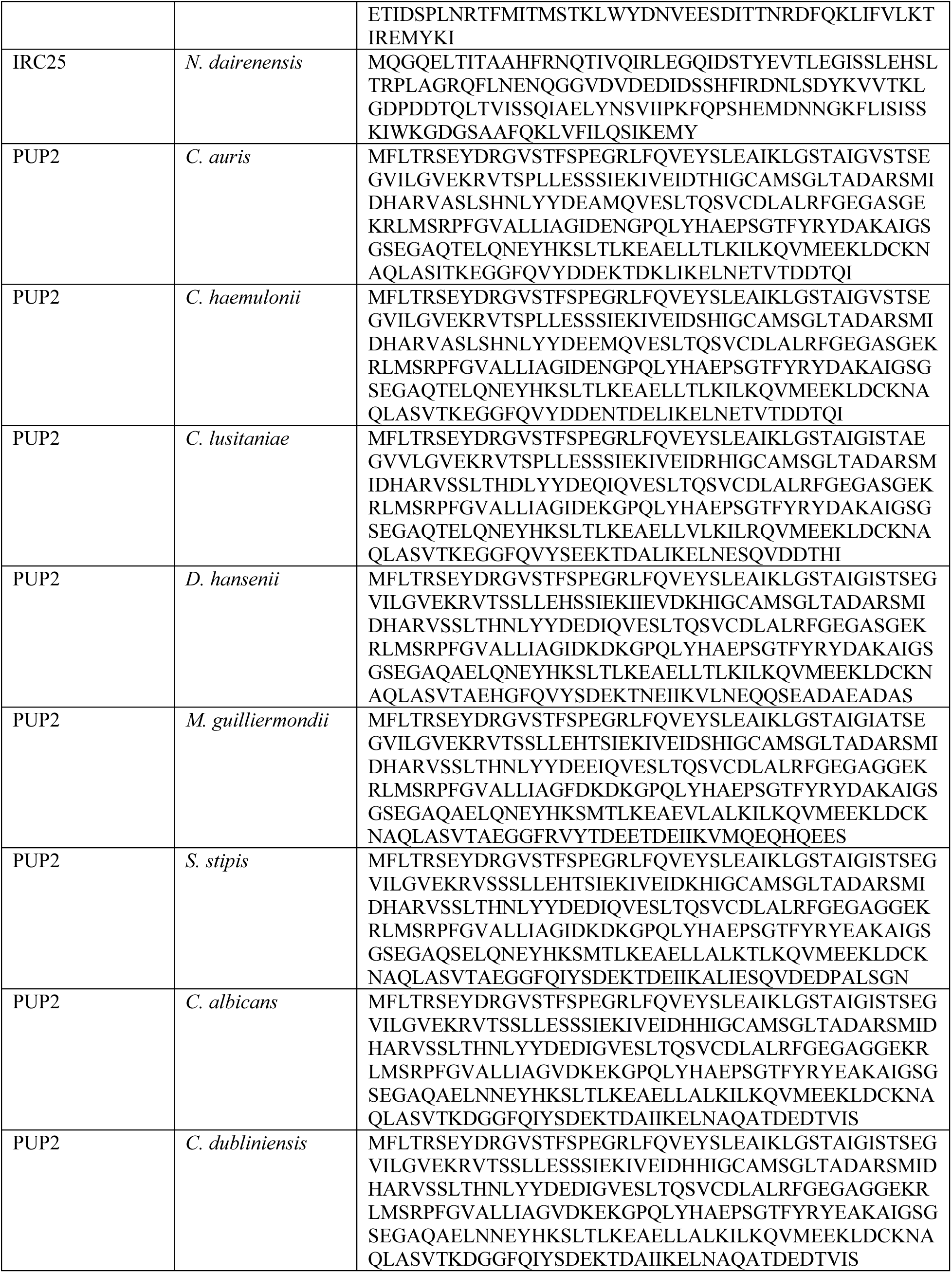
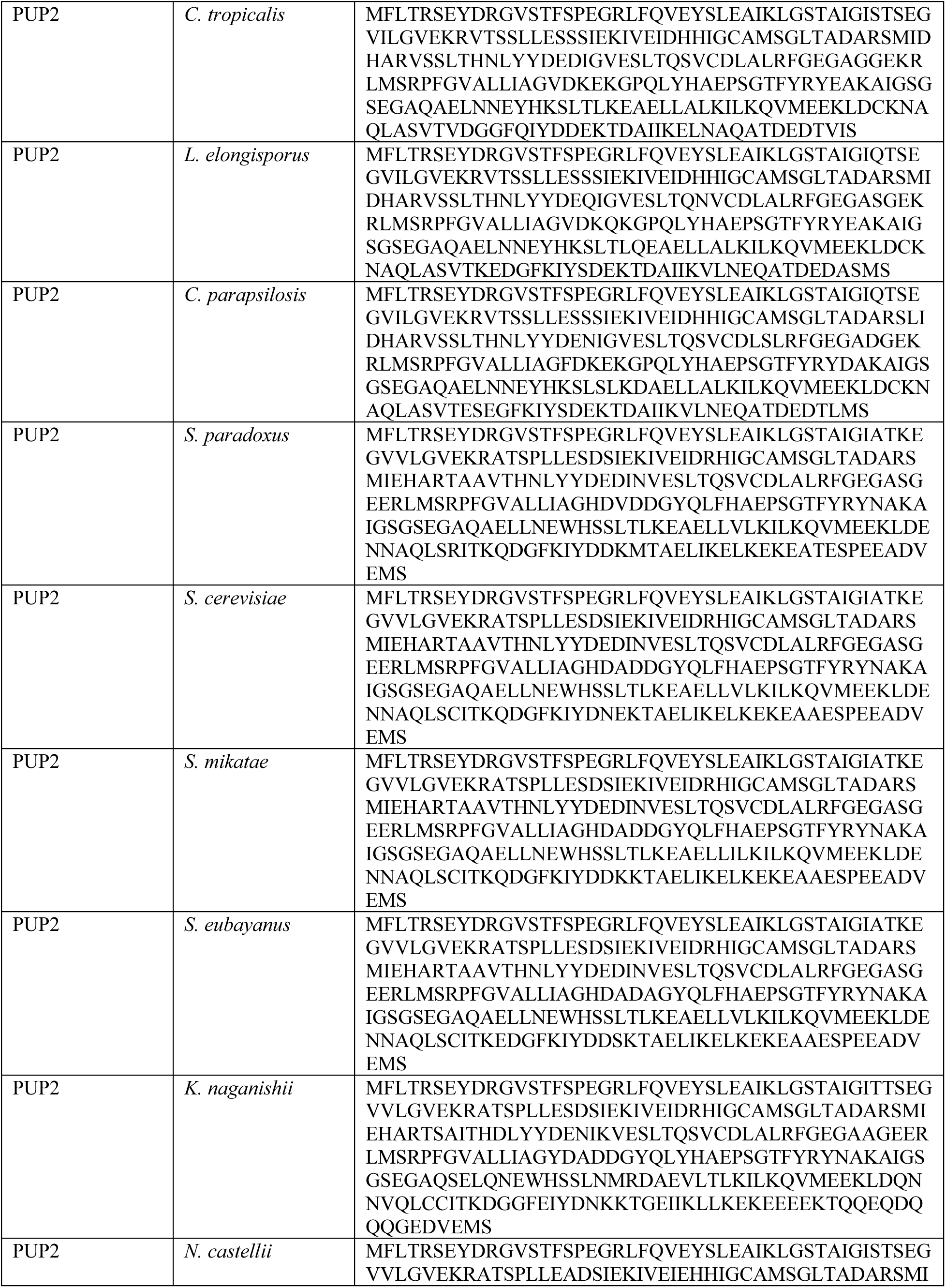
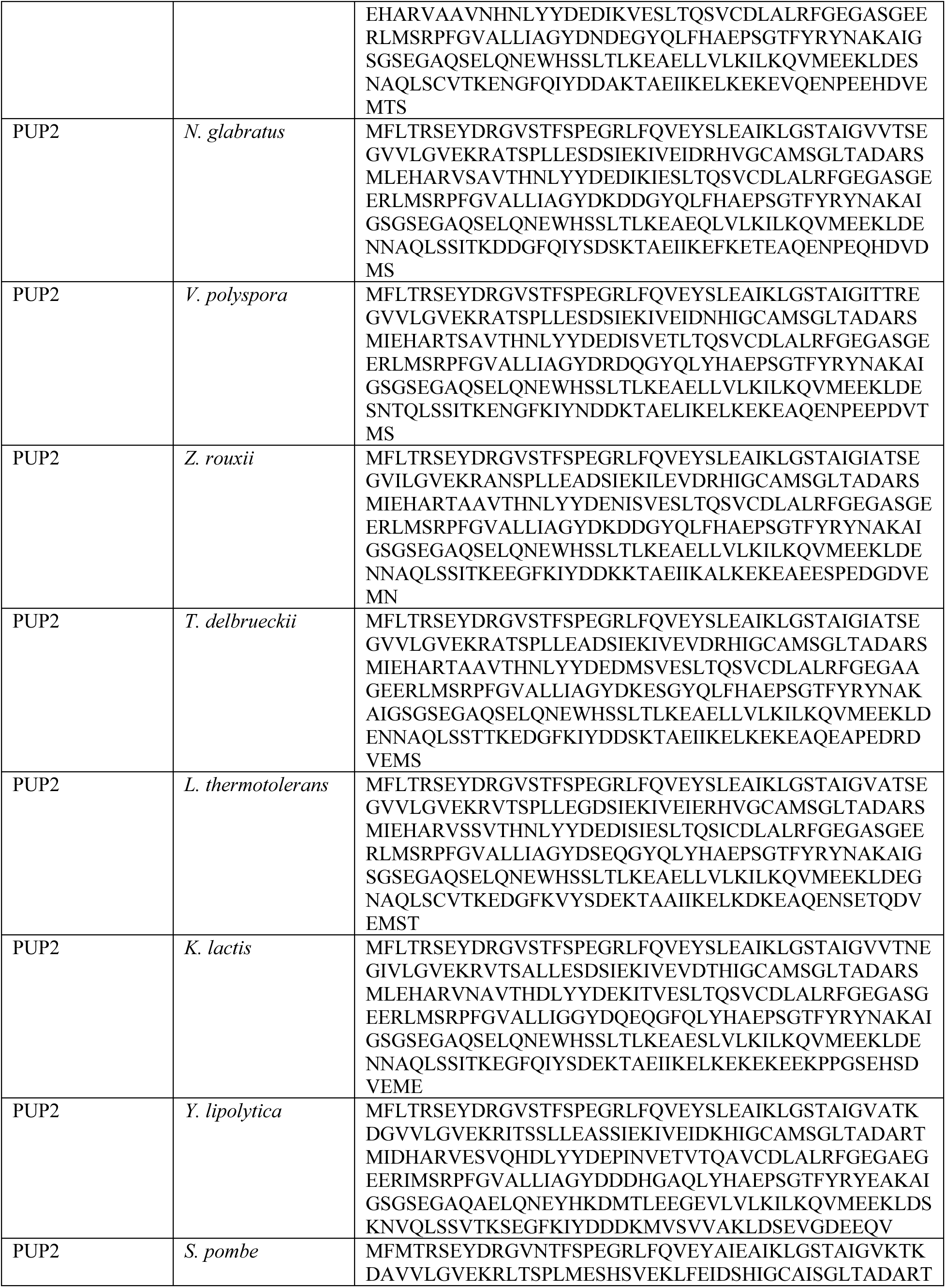
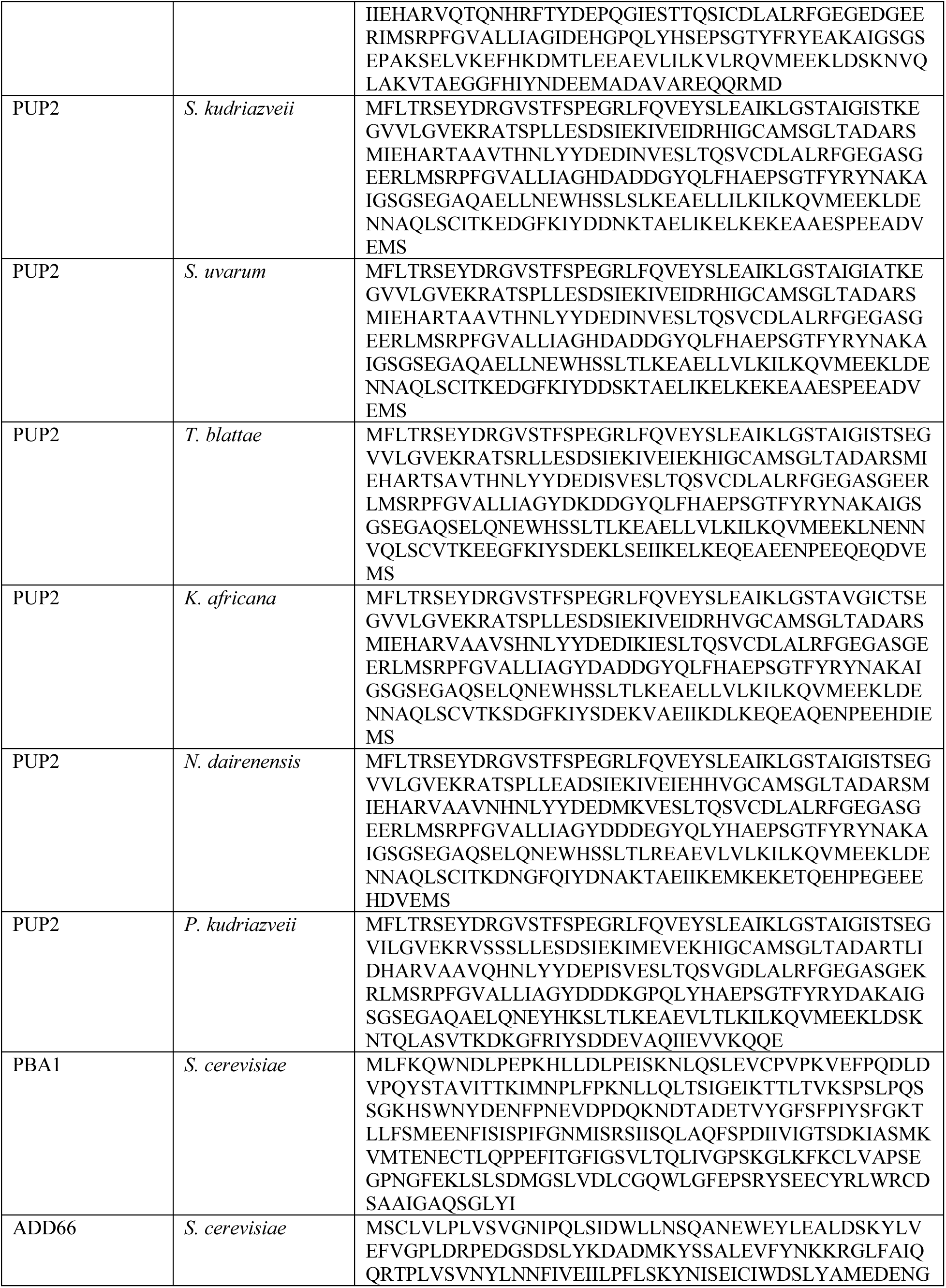
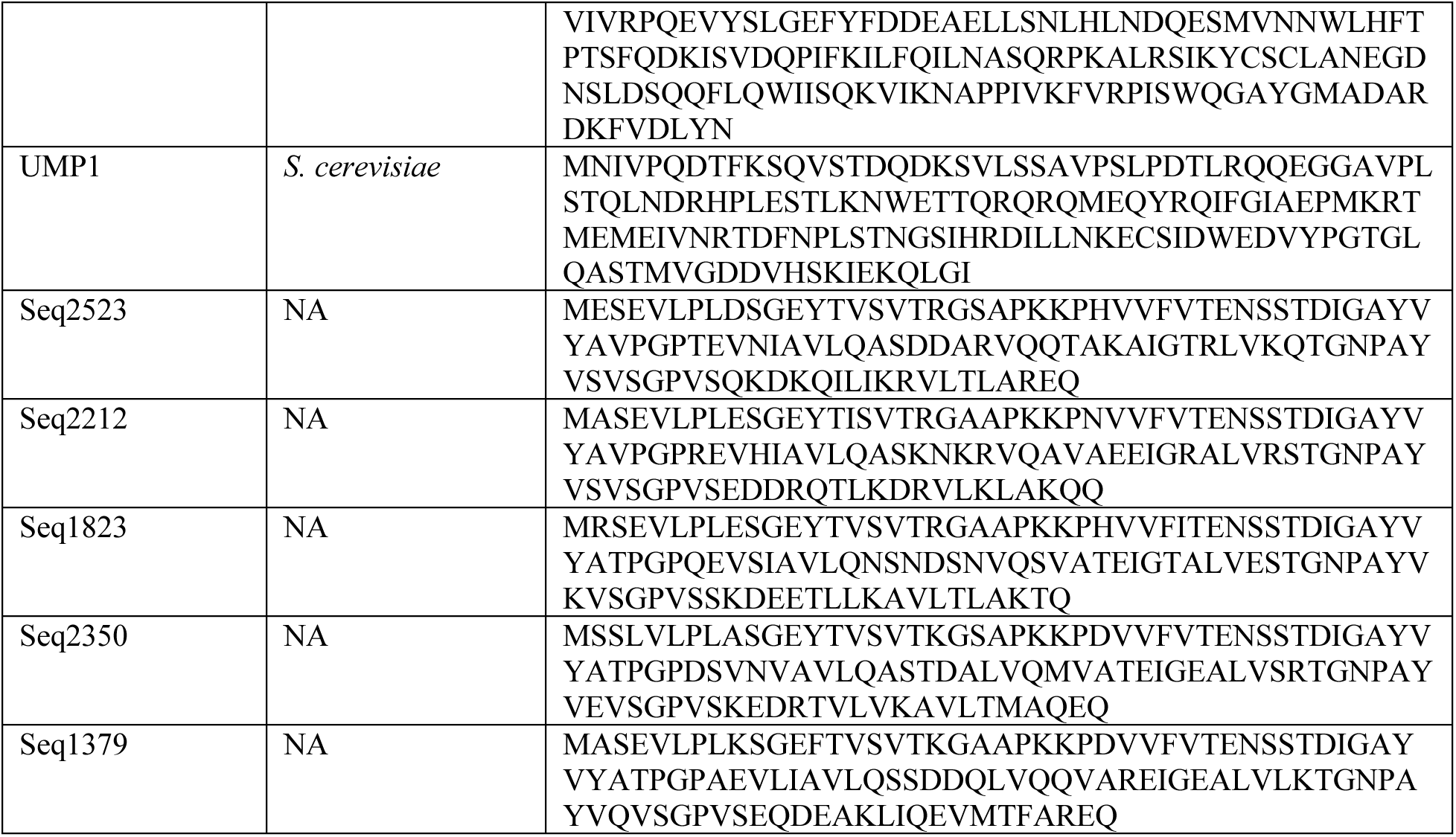
List of amino acid sequences used in this study.

**Table S5.**
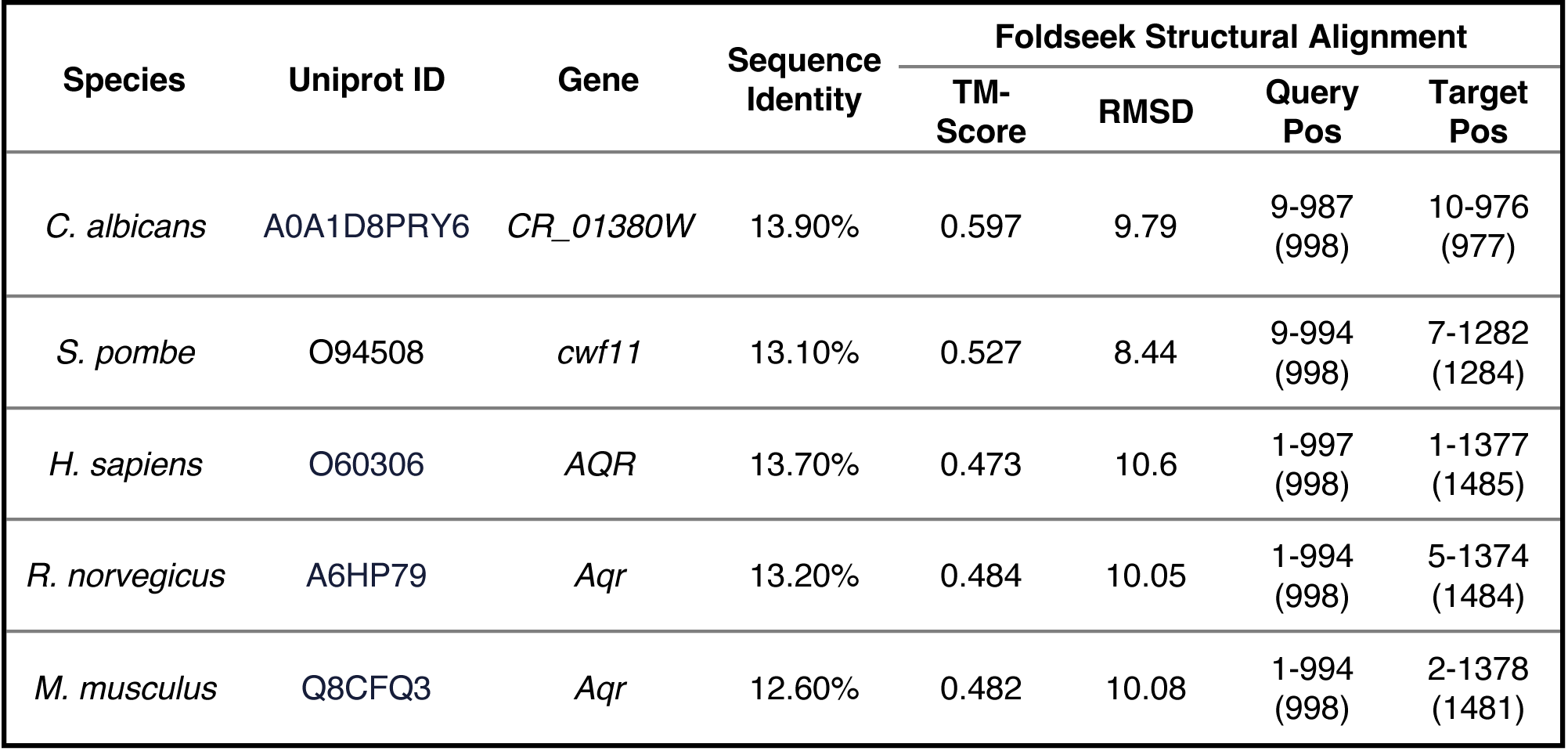
Detailed Foldseek results for B9J08_002475.

| Species | Uniprot ID | Gene | Sequence Identity | Foldseek Structural Alignment |  |  |  |
| --- | --- | --- | --- | --- | --- | --- | --- |
|  |  |  |  | TM-Score | RMSD | Query Pos | Target Pos |
| *C. albicans* | A0A1D8PRY6 | *CR_01380W* | 13.90% | 0.597 | 9.79 | 9-987 (998) | 10-976 (977) |
| *S. pombe* | O94508 | *cwf11* | 13.10% | 0.527 | 8.44 | 9-994 (998) | 7-1282 (1284) |
| *H. sapiens* | O60306 | *AQR* | 13.70% | 0.473 | 10.6 | 1-997 (998) | 1-1377 (1485) |
| *R. norvegicus* | A6HP79 | *Aqr* | 13.20% | 0.484 | 10.05 | 1-994 (998) | 5-1374 (1484) |
| *M. musculus* | Q8CFQ3 | *Aqr* | 12.60% | 0.482 | 10.08 | 1-994 (998) | 2-1378 (1481) |

**Table S6.** Detailed Foldseek results for B9J08_000767.

| Species | Uniprot ID | Gene | Sequence Identity | Foldseek Structural Alignment |  |  |  |
| --- | --- | --- | --- | --- | --- | --- | --- |
|  |  |  |  | TM-Score | RMSD | Query Pos | Target Pos |
| *S. cerevisiae* | P39011 | *BEM4* | 16.50% | 0.66 | 4.98 | 18-539<br>(543) | 33-632<br>(633) |
| *X. laevis* | O93614 | *rap1gds1-a* | 14.80% | 0.581 | 8.31 | 4-541<br>(543) | 51-604<br>(607) |
| *H. sapiens* | P52306 | *RAP1GDS1* | 14.20% | 0.582 | 8.44 | 4-541<br>(543) | 51-604<br>(607) |
| *M. musculus* | E9Q912 | *Rap1gds1* | 14.20% | 0.577 | 8.46 | 4-541<br>(543) | 51-604<br>(607) |
| *S. pombe* | O60077 | *arz1* | 13.30% | 0.467 | 11.03 | 15-543<br>(543) | 4-492<br>(492) |

**Table S7.**
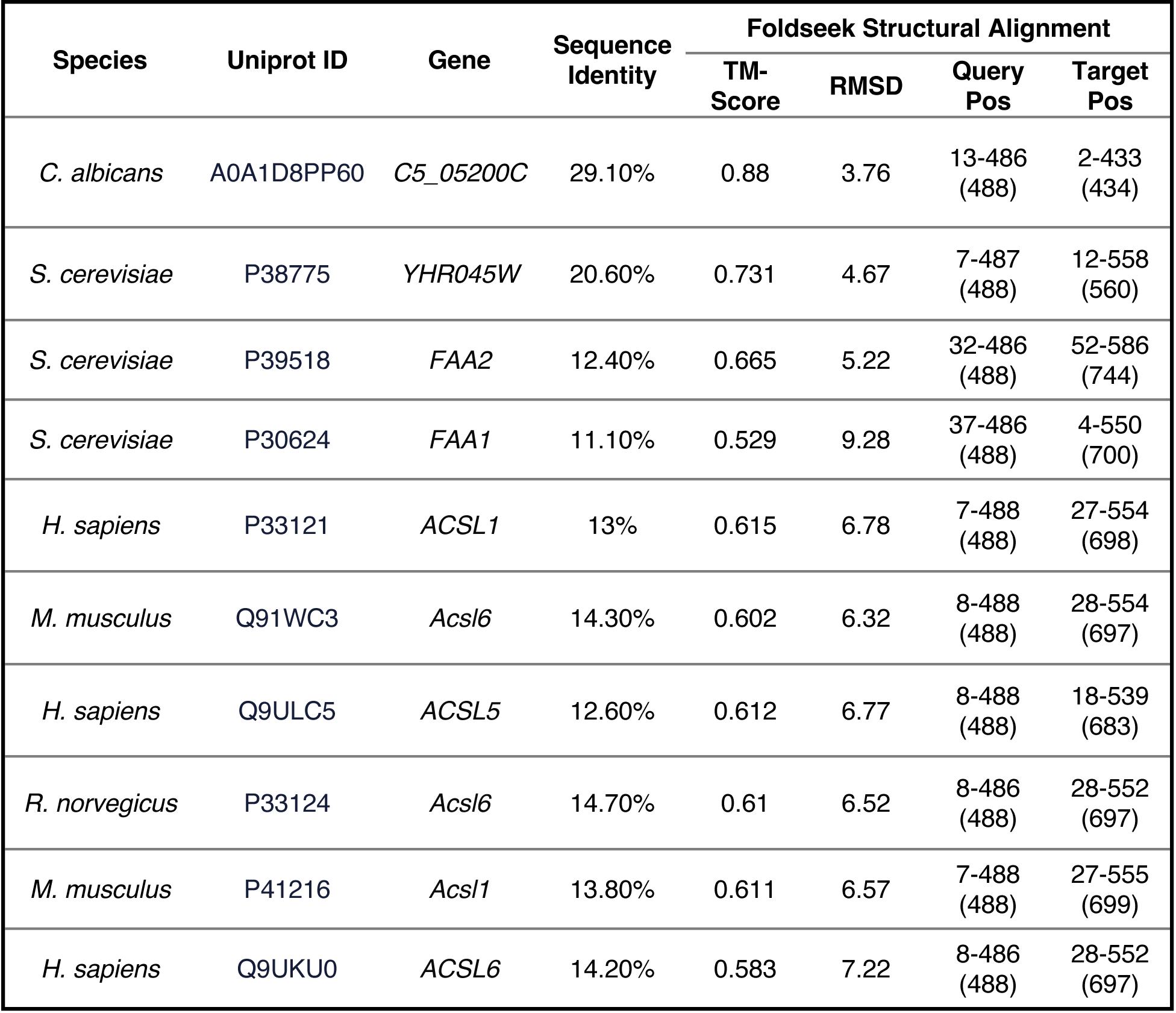
Detailed Foldseek results for B9J08_003970.

**Table S8.** Detailed Foldseek results for B9J08_000884.

| Species | Uniprot ID | Gene | Sequence Identity | FoldSeek Structural Alignment |  |  |  |
| --- | --- | --- | --- | --- | --- | --- | --- |
|  |  |  |  | TM-Score | RMS D | Query Pos | Target Pos |
| *C. albicans* | A0A1D8PTK4 | *CR_07690W* | 21.60% | 0.729 | 3.31 | 1-107 (111) | 1-104 (111) |
| *S. cerevisiae* | Q12245 | *POC4* | 20.40% | 0.551 | 4.81 | 2-107 (111) | 6-137 (148) |
| *S. pombe* | Q9P7J0 | *pac4* | 20.60% | 0.685 | 3.43 | 2-107 (111) | 7-117 (124) |
| *A. thaliana* | Q9LER5 | *PBAC3* | 13.00% | 0.691 | 2.85 | 2-107 (111) | 13-121 (124) |
| *M. musculus* | P0C7N9 | *Psmg4* | 12.60% | 0.705 | 2.91 | 2-107 (111) | 15-115 (123) |
| *H. sapiens* | Q5JS54 | *PSMG4* | 10.80% | 0.69 | 3.17 | 2-107 (111) | 15-115 (123) |

**Table S9.**
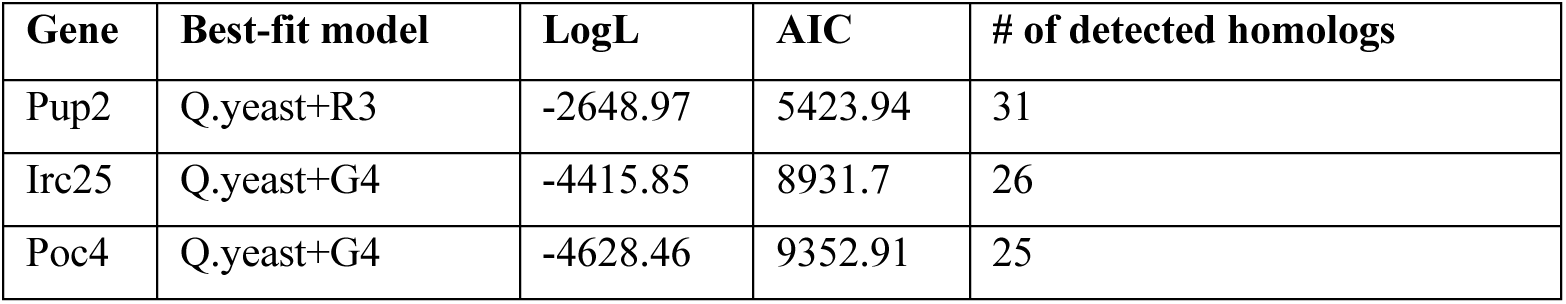
Type or paste caption here. Create a page break and paste in the table above the caption.

**Table S10.** AlphaFold 3 summary statistics for models used in this study.

| Structure | Model | pTM | ipTM | has_clash | fraction_disordered | ranking_score |
| --- | --- | --- | --- | --- | --- | --- |
| *C. auris* natural | 1 | 0.82 | 0.81 | 0 | 0.06 | 0.84 |
| *C. auris* natural | 2 | 0.81 | 0.81 | 0 | 0.06 | 0.84 |
| *C. auris* natural | 3 | 0.81 | 0.81 | 0 | 0.05 | 0.83 |
| *C. auris* natural | 4 | 0.81 | 0.8 | 0 | 0.06 | 0.83 |
| *C. auris* natural | 5 | 0.81 | 0.8 | 0 | 0.05 | 0.83 |
| *S. cerevisiae* natural | 1 | 0.79 | 0.78 | 0 | 0.07 | 0.81 |
| *S. cerevisiae* natural | 2 | 0.78 | 0.77 | 0 | 0.06 | 0.8 |
| *S. cerevisiae* natural | 3 | 0.79 | 0.78 | 0 | 0.07 | 0.81 |
| *S. cerevisiae* natural | 4 | 0.79 | 0.77 | 0 | 0.06 | 0.81 |
| *S. cerevisiae* natural | 5 | 0.78 | 0.77 | 0 | 0.06 | 0.81 |
| *C. auris* w/ ScPoc4 | 1 | 0.81 | 0.81 | 0 | 0.05 | 0.84 |
| *C. auris* w/ ScPoc4 | 2 | 0.8 | 0.8 | 0 | 0.05 | 0.82 |
| *C. auris* w/ ScPoc4 | 3 | 0.8 | 0.8 | 0 | 0.06 | 0.83 |
| *C. auris* w/ ScPoc4 | 4 | 0.8 | 0.79 | 0 | 0.05 | 0.82 |
| *C. auris* w/ ScPoc4 | 5 | 0.8 | 0.8 | 0 | 0.05 | 0.83 |
| *C. auris* w/ Seq2523 | 1 | 0.82 | 0.81 | 0 | 0.06 | 0.84 |
| *C. auris* w/ Seq2523 | 2 | 0.82 | 0.81 | 0 | 0.06 | 0.84 |
| *C. auris* w/ Seq2523 | 3 | 0.82 | 0.81 | 0 | 0.06 | 0.84 |
| *C. auris* w/ Seq2523 | 4 | 0.81 | 0.8 | 0 | 0.06 | 0.83 |
| *C. auris* w/ Seq2523 | 5 | 0.81 | 0.8 | 0 | 0.06 | 0.83 |
| *C. auris* w/ Seq2212 | 1 | 0.8 | 0.8 | 0 | 0.06 | 0.83 |
| *C. auris* w/ Seq2212 | 2 | 0.78 | 0.8 | 0 | 0.06 | 0.82 |
| *C. auris* w/ Seq2212 | 3 | 0.78 | 0.79 | 0 | 0.06 | 0.82 |
| *C. auris* w/ Seq2212 | 4 | 0.78 | 0.79 | 0 | 0.06 | 0.81 |
| *C. auris* w/ Seq2212 | 5 | 0.78 | 0.79 | 0 | 0.06 | 0.81 |
| *C. auris* w/ Seq1379 | 1 | 0.79 | 0.78 | 0 | 0.06 | 0.81 |
| *C. auris* w/ Seq1379 | 2 | 0.79 | 0.78 | 0 | 0.06 | 0.81 |
| *C. auris* w/ Seq1379 | 3 | 0.77 | 0.78 | 0 | 0.05 | 0.8 |
| *C. auris* w/ Seq1379 | 4 | 0.77 | 0.78 | 0 | 0.05 | 0.8 |
| *C. auris* w/ Seq1379 | 5 | 0.76 | 0.78 | 0 | 0.05 | 0.79 |
| *C. auris* w/ Seq2350 | 1 | 0.77 | 0.76 | 0 | 0.06 | 0.79 |
| *C. auris* w/ Seq2350 | 2 | 0.77 | 0.76 | 0 | 0.05 | 0.79 |
| *C. auris* w/ Seq2350 | 3 | 0.77 | 0.76 | 0 | 0.06 | 0.79 |
| *C. auris* w/ Seq2350 | 4 | 0.77 | 0.76 | 0 | 0.06 | 0.79 |
| *C. auris* w/ Seq2350 | 5 | 0.77 | 0.76 | 0 | 0.05 | 0.79 |
| *C. auris* w/ Seq1823 | 1 | 0.79 | 0.79 | 0 | 0.07 | 0.82 |
| *C. auris* w/ Seq1823 | 2 | 0.8 | 0.79 | 0 | 0.06 | 0.82 |
| *C. auris* w/ Seq1823 | 3 | 0.79 | 0.79 | 0 | 0.06 | 0.82 |
| *C. auris* w/ Seq1823 | 4 | 0.78 | 0.79 | 0 | 0.05 | 0.81 |
| *C. auris* w/ Seq1823 | 5 | 0.77 | 0.78 | 0 | 0.07 | 0.81 |

**Data S1. (separate file)**

Type or paste caption here.

